# Stress pathway outputs are encoded by pH-dependent phase separation of its components

**DOI:** 10.1101/2024.02.24.581896

**Authors:** Yuliia Didan, Milad Ghomlaghi, Lan K. Nguyen, Dominic C. H. Ng

## Abstract

Signal processing by intracellular kinases control near all biological processes but how precise functions of signal pathways evolve with changed cellular contexts is poorly understood. Functional specificity of c-Jun N-terminal Kinases (JNK) activated in response to a broad range of pathological and physiological stimuli are partly encoded by signal strength. Here we reveal that intracellular pH (pHi) is a significant component of the JNK regulatory network and defines JNK signal response to precise stimuli. We showed that nuanced fluctuations in physiological pHi regulates JNK activity in response to cell stress. Interestingly, the relationship between pHi and JNK activity was dependent on specific stimuli and upstream kinases involved in pathway activation. Cytosolic alkalinisation promoted phase transition of upstream ASK1 to augment JNK activation. While increased pHi similarly induced JNK2 to form condensates, this led to attenuated JNK activity. Mathematical modelling of feedback signalling incorporating pHi and differential contribution by JNK2 and ASK1 condensates was sufficient to delineate the strength of JNK signal response to specific stimuli. This new knowledge of pHi regulation with consideration of JNK2 and ASK1 contribution to signal transduction may delineate oncogenic versus tumour suppressive functions of the JNK pathway and cancer cell drug responses.

## Introduction

Signal transduction by intracellular kinases integrates and process biochemical information generated by a multitude of cell intrinsic and extrinsic stimuli. This is required to co-ordinate highly specific cell responses to their changing environments. The c-Jun N-terminal kinases (JNK) are conserved members of the mitogen-activated protein kinase (MAPK) family, characterized by activation through a tiered phosphorylation cascade (MAP3K→ MAP2K → MAPK) and serve as central nodes within these signal regulatory networks ^1^. The JNK pathway is activated by diverse sets of stimuli, including growth factors, cytokines, proteotoxic, environmental and metabolic stress ^2^. Following activation, JNK promotes a wide range of cellular responses, including cell fates that are often considered mutually exclusive, such as survival versus programmed death, and proliferation versus differentiation ^2^. In cancer for example, both oncogenic and tumour suppressor roles for JNK signals are well recognized ^3,4^. The kinetics of JNK activation is a key mechanism that delineates between seemingly contradictory functions of the pathway. Sustained activation of JNK leads to cell cycle arrest and programmed death, while transient activation of the pathway is associated with survival and proliferation ^5^. This raises the fundamental question of how JNK activation is changed in different cell contexts.

As the terminal kinases within a tiered cascade, JNK MAPKs receive signals from multiple upstream MAP3K activated by specific stimuli. For example, MLK3 primarily signals to JNK following activation by the inflammatory cytokine, TNFα ^6^, while ASK1 is preferentially activated by oxidative stress ^7^ and ASK3 responds to hyperosmolarity ^8^. The signal transduction mediated by distinct MAP3Ks may result in differential engagement of feedback and feedforward loops, ultimately defining dynamic JNK signals and downstream cellular responses to stimulation. The organization of signal proteins within the cytosolic environment is also a critical determinant of how signals are propagated through pathways. An emerging concept is that signal networks may further evolve with intracellular pH (pHi) due to altered protonation state and regulation of higher-ordered signalling complexes ^9,10^.

Intracellular pH (pHi) is kept tightly regulated within a narrow range under physiological conditions to guard against excessive acidification or alkalinization of the cytosol that would be detrimental to protein function ^11^. However, it is now evident that small fluctuations in pHi regulate fundamental processes ^12–14^ and changes in pHi further reflects pathological cell states ^15^. Notably, an increasing body of evidence supports a direct role for pHi in regulating signal transduction. pHi influences the stability of β-catenin and subsequent activation of the Wnt pathway ^10^. Similarly, pHi-dependent conformational changes in Gα subunits are important determinants of G-protein coupled receptor-initiated signalling ^16^. Fluctuations in pHi accompany the programmed death and proliferation of cancer cells, with JNK signalling being an important regulator of both processes. Given that phosphorylation networks regulate ion channels and transporters that determine cytosolic pH ^17,18^, pHi may represent embedded components of kinase regulatory networks that warrant more in-depth consideration. Moreover, the cross-regulation between protein and proton [H^+^] components within signalling networks are largely unexplored.

Here, we utilized live-cell biosensor measurements to reveal a tight association between JNK activity and changes in pHi in response to stress stimulation. We demonstrated that pHi was sufficient to regulate JNK signal output, and this was mediated through pH-dependent phase transition and altered signal transduction of tiered kinases embedded within the JNK pathway. Interestingly, pH-regulated ASK1 at the MAP3K tier contribute differently to signal transduction compared to pH-dependent phase transitions of JNK. Our mathematical model, which incorporates signal feedback on JNK activation, pHi, and the differential contribution by kinase condensates, accurately predicted JNK pathway responses to specific stimuli and explained how different JNK signal strengths are generated. Our study helps delineate the contextual functions of JNK signals, such as in drug-resistant cancers where JNK has been reported to serve dual functions.

## Results

### Cytosolic pH influences stress-stimulated JNK signalling

An analysis of the Genomics of Drug Sensitivity in Cancer (GDSC) database indicated that cancer cell type sensitivity to JNK-targeting drugs was consistently negatively correlated with pHi (Fig. S1A). There may be several reasons underlying this association, but our observations suggest a potential link between JNK signal transduction and pHi. This prompted us to investigate this more closely through live-cell measurements of pHi and JNK activity. Our studies utilized a fluorogenic dye (pHrodo) and a protein-based ratiometric sensor (ClopHensor) as two independent sensors of pHi in live cells, in conjunction with a JNK activity reporter (JNK-KTR ^19,20^). Our measures of pHrodo fluorescence and ClopHsensor green to cyan ratios (Fig. S1, B to F) were carefully calibrated against internal pH standards to enable direct comparison of pHi in different cell contexts and between sensors. As JNK is typically associated with stress responses, we first evaluated pHi and JNK activity responses to proinflammatory TNFα as a protypical stress activator of the pathway ^21^. TNFα treatment of HEK293 cells led to JNK activation and modulation of pHi (Fig. 1A). Specifically, pHi decreased as JNK activity rapidly increased following TNFα treatment (Fig. 1B). After reaching peak JNK activation at around 25 mins, JNK activity decreased while pHi increased over time (Fig. 1B). Thus, JNK activity was inversely associated with changes in pHi following TNFα stimulation (Fig. 1C). In an expanded analysis of a panel of immortalized cell lines with measured pHi ranging between 6.9 to 7.6 (Fig. S1, C to F), we observed that the dynamics of JNK activation in response to TNFα varied markedly between different cell types (Fig. 1D). TNFα-stimulated JNK signals appeared to be enhanced in cells with comparatively acidic pHi and dampened in cells with higher pHi (Fig. 1E). Therefore, the strength of JNK signals generated by TNFα were negatively correlated with pHi both between different cell types and within the same cells over time.

**Fig. 1.**
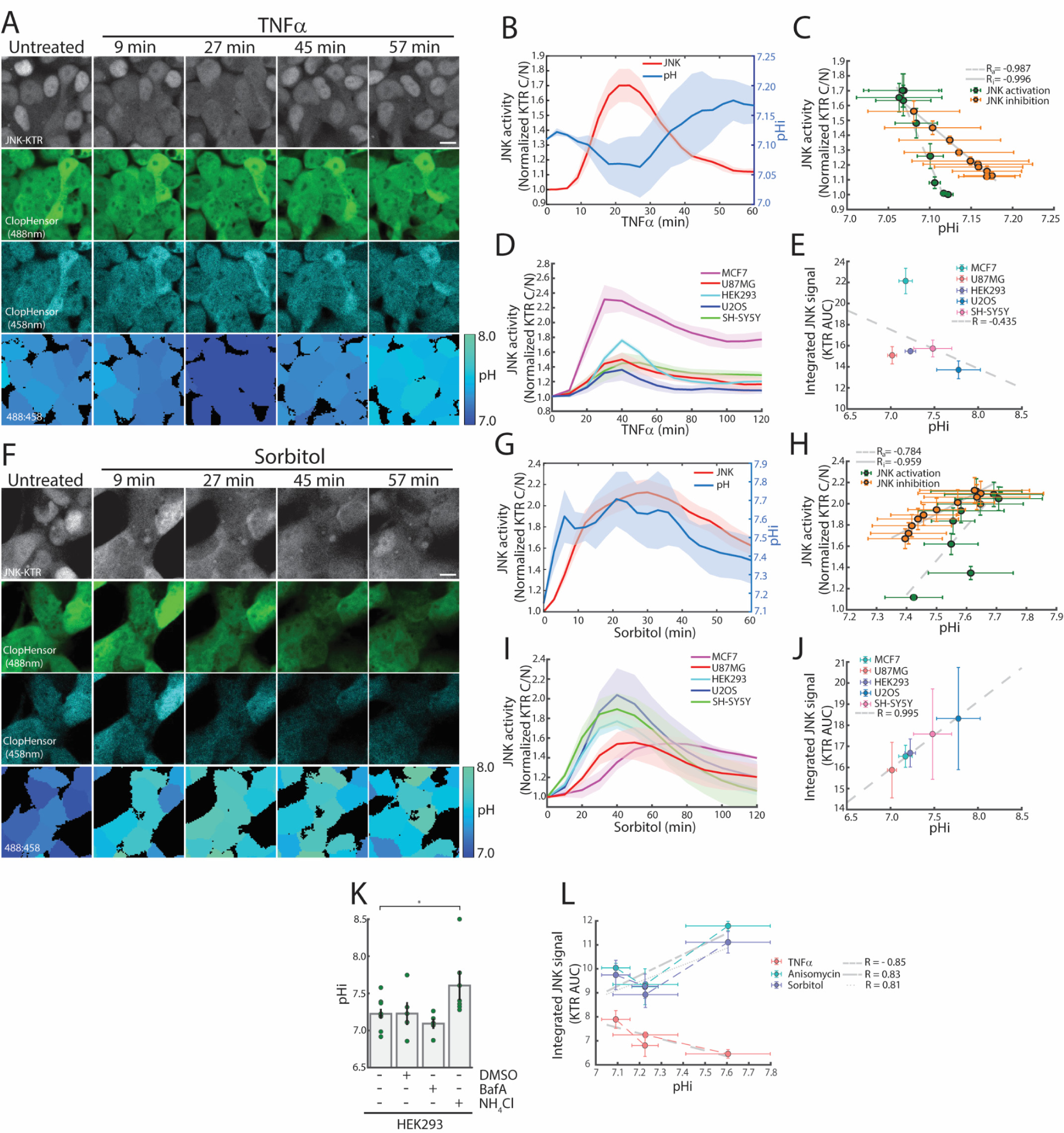
Intracellular pH (pHi) modifies stress-stimulated activation of JNK MAPK. (**A**) Representative images of JNK activation (KTR, where JNK activation is correlated with nuclear export of the sensor) and changed pHi indicated by a ratio of ClopHensor green to cyan fluorescence in TNFα (10 ng/ml) stimulated HEK293 cells. (**B**) JNK activity (KTR) and pHi responses to time-course treatment with TNFα (n = 5). (**C**) JNK activity during activation and inactivation response phase to TNFα stimulation was correlated to pHi determined with ClopHensor. (**D**) TNFα (10 ng/ml) time course treatment in a panel of cell lines (U87MG, MCF7, HEK293, SH-SY5Y and U2OS), n ≥ 3 independent experiments. (**E**) Intracellular pHi across cell types negatively correlated with TNFα-stimulated JNK signal capacity measured as integrated area under JNK-KTR C/N curve. (**F**) Representative images of JNK activity and pHi in sorbitol (150 mM) stimulated HEK293 cells (**G**) Quantified measure of JNK activity and pHi response to a time course of sorbitol (150 mM) treatment (n = 5). (**H**) JNK activation and inactivation following sorbitol treatment correlated with pHi. (**I**) JNK activity response to a sorbitol (150 mM) time course across panel of cell lines, n ≥ 3 independent experiments. (**J**) Intracellular pHi across cell types were positively correlated with sorbitol-stimulated JNK signal capacity (integrated area under JNK-KTR C/N curve). (**K**) Cells treated with ammonium salt (50 mM NH_4_Cl) or bafilomycin A1 (50 µM BafA) and pHi determined with ClopHensor (n = 8, per condition). (**L**) Correlation between JNK activity and pHi in HEK293 cells pretreated with NH_4_Cl, BafA or vehicle control followed by stress stimulation. Values represent mean±SEM. “R” indicates Pearson correlation coefficients between pHi and JNK activity including correlation coefficients between pHi and JNK activation (R_a_) or inactivation (R_i_). Unpaired t test, *p < 0.05 Scale bars, 10 µm.

We next investigated whether pHi was similarly associated with JNK activity stimulated by hyperosmolarity, a physical stressor and robust activator of the JNK pathway ^22^. Surprisingly, we found that pHi fluctuations were closely associated with JNK activity in HEK293 cells exposed to hyperosmolarity (Fig. 1F). In response to osmotic stress, pHi increased alongside JNK activation, peaked and then subsequently decreased as JNK signals were deactivated (Fig. 1, G and H). Similarly, we observed a strong positive correlation between cell type specific pHi and JNK signal strength stimulated by hyperosmolarity (Fig. 1, I and J). In sharp contrast to TNFα and osmotic stress, we did not observe a clear correlation between time or cell type-dependent pHi with JNK activity stimulated by the ribotoxic stressor, anisomycin (Fig S1, G to J). Thus, stress-stimulated JNK signalling was closely associated with pHi, although the nature of the association appeared to be stimulus dependent.

To determine if a physiological change in pHi was sufficient to alter JNK activity, we utilized bafilomycin A1 (BafA), a potent inhibitor of V-ATPases, to inhibit acidification of lysosomes leading to increased cytosolic [H^+^] ^23^, and ammonium chloride (NH_4_Cl) that causes permeation of cells with weak NH3^+^ base to reduce cytosolic [H^+^] ^11^. We treated HEK293 cells, which had pHi in the middle of the physiological pHi range and confirmed that NH_4_Cl treatment led to cytosolic alkalinization or increased pHi while BafA resulted in acidification or reduced pHi (Fig. 1K). We then assessed the effect of chemically modulated pHi on JNK activity in response to our three candidate stress stimuli (Fig. S1K). When JNK activity was plotted against pHi from all treatments, we found that acidification and alkalinization of the cytosol led to a predictable shift in JNK signalling capacity, which reinforced a strong inverse relationship between pHi and TNFα induced JNK activity and an opposite relationship for anisomycin and sorbitol (Fig. 1L). Our studies demonstrate that pHi contributes to determining the strength of JNK activation following stimulation. Given that JNK regulates membrane [H^+^] transporters ^24^, JNK activation may promote further changes in pHi, inducing a complex interplay that is dependent on the stress signal context.

### Stress-stimulated clustering of JNK pathway kinases is regulated by pHi

As pH alters the charge states of proteins and their interactions within the crowded cytosol ^25^, we investigated the subcellular organisation of JNK pathway signal components to gain insights into stress-responsive and pH-regulated signalling. We focused our attention on JNK2 and ASK1 as signal proteins in the JNK pathway reported to assemble higher ordered structures to regulate signalling ^26,27^. While ASK1 is implicated in transducing a broad range of stress signals, we found that pretreatment of HEK293 cells with an ASK1 specific inhibitor attenuated sorbitol and, to a lesser extent, anisomycin-stimulated JNK activity but did not change JNK activation in response to TNFα (Fig. S2, A and B). Similarly, ectopic expression of JNK2 enhanced JNK-KTR measured activity induced by all stress stimuli tested, while exogenous expression of ASK1 augmented sorbitol and anisomycin activation of JNK but did not change JNK responses to TNFα (Fig. S2C). This indicates that ASK1 transduces stress signals to JNK in response to hyperosmolarity and ribotoxic stress but is less involved in TNFα-stimulated responses.

Our live tracking of mCherry-tagged proteins in HEK293 cells revealed that a proportion of JNK2 and ASK1 formed visible puncta or foci in the cytosol prior to stress (Fig. 2A). In response to each stress treatment tested, the number of JNK2 foci were reduced as JNK activity (KTR) increased and the number of JNK2 foci increased as JNK activity decreased following peak activation (Fig. 2, B to D). Similarly to JNK2, the number of ASK1 foci were inversely associated with JNK activation state following TNFα stimulation (Fig. 2, E and F). Interestingly, in response to anisomycin or sorbitol, ASK1 foci numbers increased and decreased in alignment with JNK activation and deactivation, respectively (Fig. 2, G and H). Thus, while stress induced assembly of JNK2 into cytosolic foci was negatively correlated with JNK activity, the formation of ASK1 puncta were positively correlated with JNK activity in hyperosomotic and ribotoxic stress contexts that signal through ASK1 (Fig. 2I). We next compared the propensity for exogenously expressed mCherry-tagged JNK2 and ASK1 to form cytoplasmic foci in our panel of cell lines with differing pHi. The expression levels of JNK2 and ASK1, as determined by fluorescence intensity, was not a strong predictor of the propensity to form cytoplasmic foci in unstimulated cells (Fig. S2, D to H). Instead, we observed increased numbers of JNK2 and ASK1 foci in unstimulated cells with comparatively more alkaline cytosols (eg. U2OS), with the number of foci strongly correlated positively with pHi (Fig. 3, A to D).

**Fig. 2.**
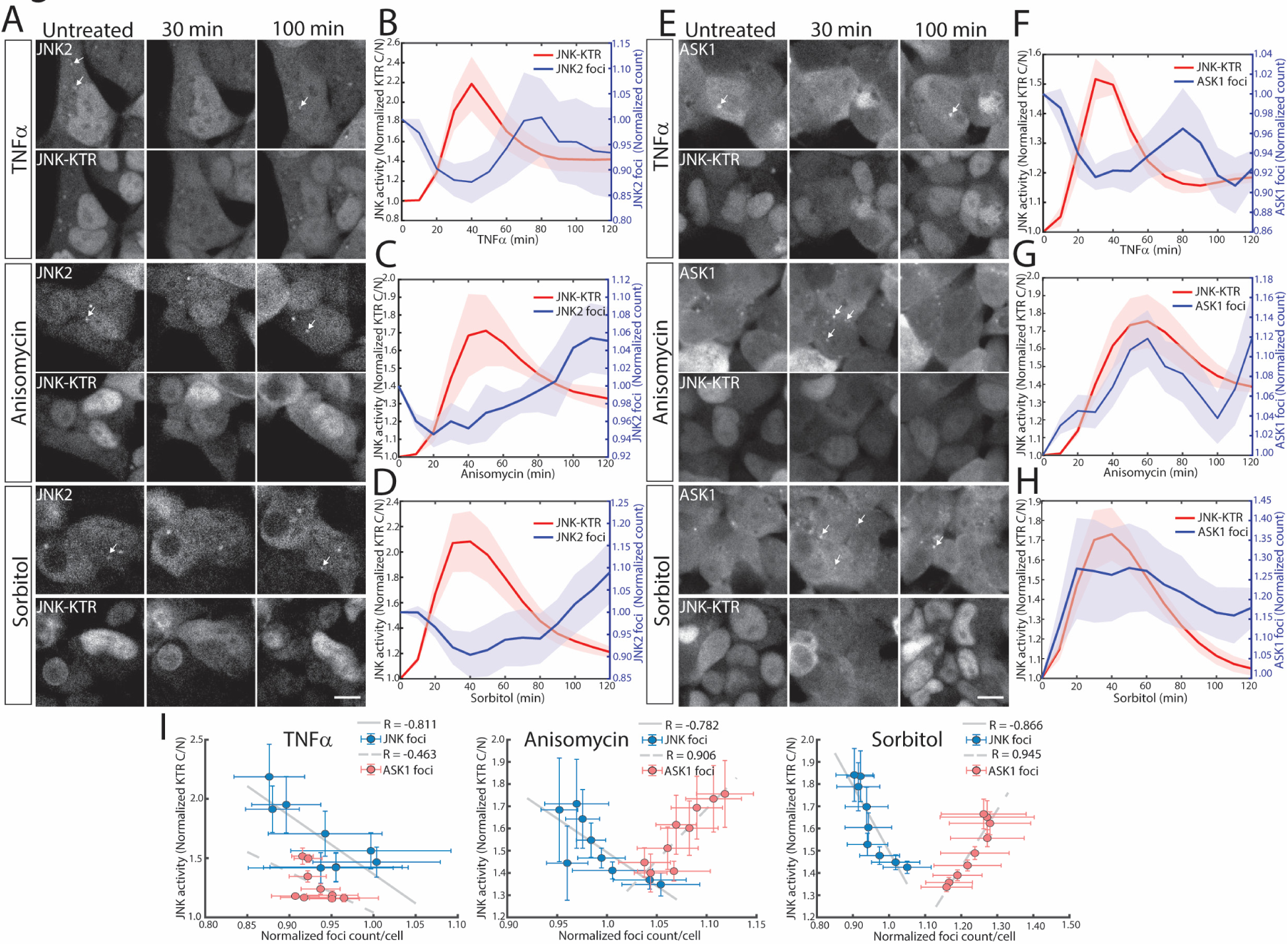
Stress-stimulated assembly of JNK2 and ASK1 cytoplasmic foci. (**A**) Representative images of HEK293 cells expressing mCherry-tagged JNK2 and JNK-KTR were treated with TNFα (10 ng/ml), anisomycin (10 ng/ml) or sorbitol (150 mM) for indicated time points. (**B** to **D**) JNK2 foci counts and JNK activity (KTR) over time course of **B**, TNFα (10 ng/ml, n = 4), **C**, anisomycin (10 ng/ml, n = 5) or **D**, sorbitol (150 mM, n = 5) treatment. (**E**) Representative images of HEK293 cells expressing mCherry-tagged ASK1 and JNK-KTR were treated with TNFα (10 ng/ml), anisomycin (10 ng/ml) or sorbitol (150 mM) for indicated time points. (**F** to **H**) ASK1 foci counts and JNK activity (KTR) over time course of **F**, TNFα (10 ng/ml, n = 7), **G**, anisomycin (10 ng/ml, n = 6) or **H**, sorbitol (150 mM, n =5) treatment. (**I**) ASK1 or JNK2 cluster counts from HEK293 cells treated with TNFα (10 ng/ml), anisomycin (10 ng/ml) or sorbitol (150 mM) were correlated with JNK activity (KTR). Values represent mean±SEM. ‘R’ values indicate Pearson correlation coefficients between JNK2 or ASK1 clusters counts and JNK activity. Scale bars, 10 µm.

**Fig. 3.**
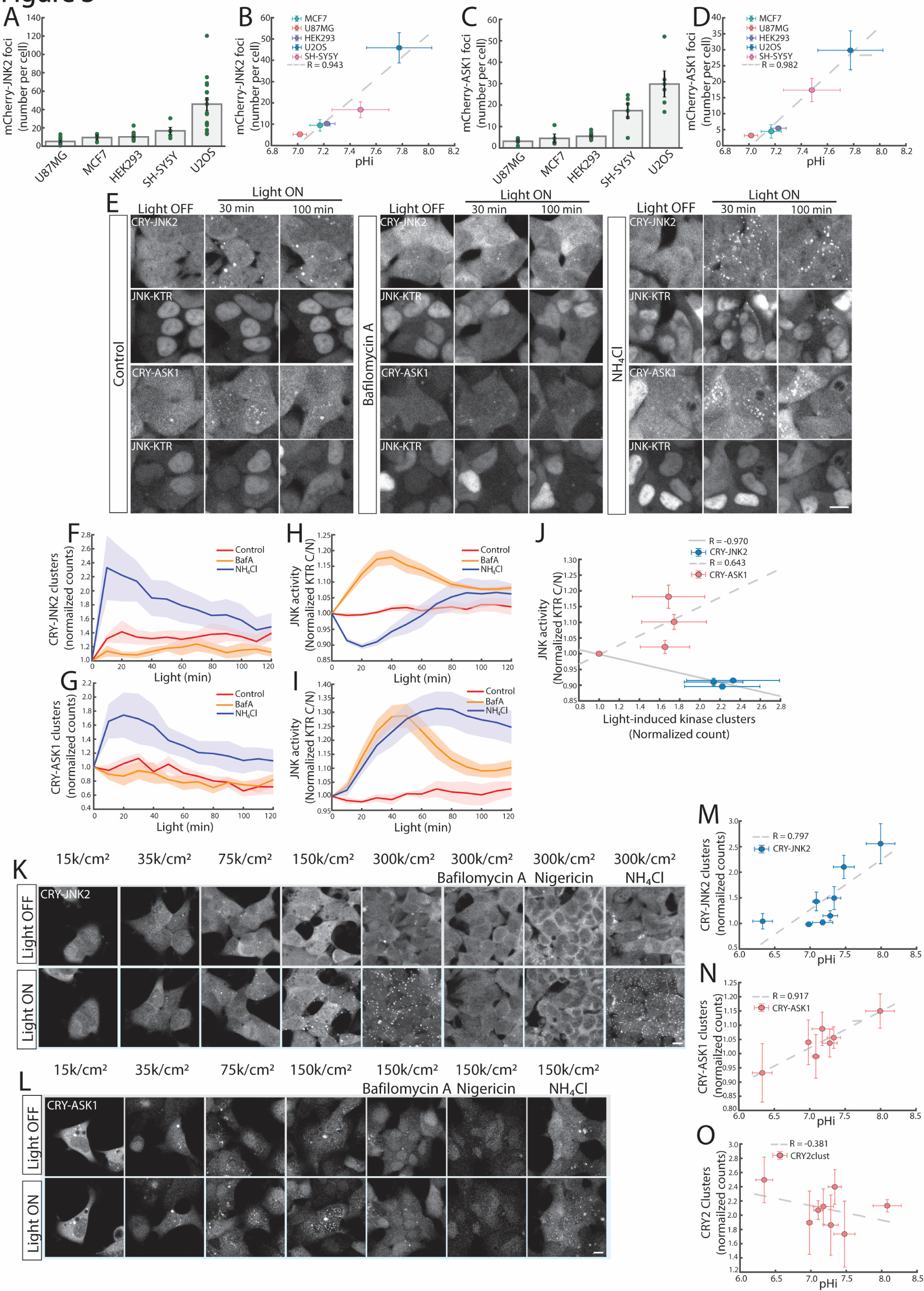
pHi regulates JNK2 and ASK1 subcellular organization and signal transduction. (**A**) mCherry-tagged JNK2 foci numbers differ by cell type, n ≥ 3 independent experiments. (**B**) JNK2 foci were positively correlated with pHi across a panel of cell lines. (**C**) mCherry-ASK1 foci numbers in different cell lines, n ≥ 3 independent experiments. (**D**) ASK1 foci were positively correlated with pHi across a cell line panel. (**E**) HEK293 cells stably expressing CRY-JNK2 or CRY-ASK1 were pretreated with NH_4_Cl (50 mM), Bafilomycin A1 (50 µM) or a vehicle control before blue-light stimulation for indicated time durations to induce kinase clustering. (**F**) Quantification of CRY-JNK2 or (**G**) CRY-ASK1 cluster formation in response to light in presence of NH_4_Cl (50 mM), Bafilomycin A1 (50 µM) or a vehicle control (n = 5, per condition). (**H**) JNK activity (KTR) in HEK293 cells expressing CRY-JNK2 or (**I**) CRY-ASK1 in response to light in presence of NH_4_Cl (50 mM), Bafilomycin A1 (50 µM) or a vehicle control (n = 5, per condition). (**J**) Correlation between light-induced clusters of CRY-JNK2 and CRY-ASK1 and JNK activity. (**K**) CRY-JNK2 or (**L**) CRY-ASK1 in HEK293 cells seeded at the indicated cell densities were pretreated with Bafilomycin A1 (50 µM), nigericin (10 µM to equilibrate to pH 7.0), NH_4_Cl (50 mM) or left untreated and light stimulated (Light On, 30 min), n ≥ 4, per condition. Correlation between (**M**) CRY-JNK2, (**N**) CRY-ASK1 and (**O**) Cry2_clust_ cluster counts and pHi determined under experimental conditions of varied cell densities or pretreatment with pH-modifying chemical treatments. Values represent mean±SEM. ‘R’ indicates Pearson correlation coefficient between cluster/foci counts and pHi or JNK activity. Scale bars, 10 µm.

To determine whether formation of JNK2 and ASK1 foci altered signal transduction and was regulated by pHi, we utilized Cry2_clust_ ^28^ to generate fusions with JNK2 and ASK1, enabling the blue light induction of clustering of kinases independently of stress stimulation. Blue light illumination of HEK293 cells induced clustering of CRY-ASK1 and CRY-JNK2 (Fig. 3E). CRY-JNK2 clusters were largely cytoplasmic while a proportion of CRY-ASK1 clusters were atypically nuclear (Fig. 3E). Light-induced JNK2 and ASK1 clustering was modestly attenuated or unaltered with BafA pretreatment but potently enhanced with NH_4_Cl (Fig. 3, F to G). Moreover, light-induced JNK activity (KTR) in cells expressing CRY-JNK2 were enhanced with BafA and was markedly attenuated with NH_4_Cl (Fig. 3H). In CRY-ASK1 cells, light-stimulated JNK signal activity was augmented with both BafA and NH_4_Cl pre-treatment (Fig. 3I). Given light-induced kinase clustering was not substantially altered by BafA, the increased activation of JNK in this context may not be specifically attributed to changes in kinase subcellular organization. In contrast, our studies indicate that increased pHi markedly enhanced kinase clustering. While light-stimulated condensate formation enhanced JNK signal activation in cells expressing CRY-ASK1, the inverse is true in cells expressing CRY-JNK2 (Fig. 3J). These findings suggest that clustering of ASK1 potentiates JNK activation while assembly of JNK2 clusters negatively correlates with JNK activity.

The formation of dense monolayers also leads to increased pHi due to regulation by cell surface adhesive receptors ^29^. We confirmed increased cytoplasmic and nuclear pH with increased seeding density of HEK293 cells (Fig. S3, A and B). The increased cell density led to enhanced clustering of CRY-JNK2 in response to light (Fig. 3K and Fig. S3C). The light-induced formation of JNK2 clusters within dense monolayers was further promoted with NH_4_Cl alkalinization (Fig. 3K). In contrast, acidification with BafA or equilibration of pHi with extracellular pH (7.0) using nigericin, a carboxylic ionophore that promotes K^+^/H^+^ exchange ^30^, attenuated CRY-JNK2 cluster formation (Fig. 3K and Fig. S3D). Unlike JNK2, light induction of CRY-ASK1 clusters was not substantially increased at higher cell densities (Fig. 3L and Fig. S3E) but was inhibited with BafA or nigericin and augmented with NH_4_Cl (Fig. 3L, and Fig. S3F). Thus, the increased pH from high density cell cultures may not be sufficient to augment ASK1 foci formation compared to chemical alkalinization which results in higher pH values (Fig. S3, A and B). As a control, the light-responsive oligomerization of the Cry2_clust_ domain alone was not enhanced by culturing cells at higher densities and not substantially altered through chemical modulation of pHi (Fig. S3, G to I). Thus, JNK2 and ASK1 show a specific sensitivity to the cytosolic pH environment. A plot of cluster counts against measured pHi at various cell densities and chemical treatments indicates a strong positive association between pHi and the assembly of JNK2 and ASK1 clusters and confirms that Cry2_clust_ was not strongly correlated with pHi (Fig. 3, M to O). Thus, there is substantial pHi influence over the subcellular organization of JNK signal components. The formation of ASK1 and JNK2 cytosolic foci is enhanced under alkaline pHi and this was associated with potentiated or attenuated JNK activation respectively, indicating differential effects on signal flux through the pathway.

### pH-regulated phase transition of JNK pathway kinases

The phase transition of kinases and self-assembly into protein condensates is sensitive to pH fluctuations and increasingly implicated in signal transduction ^31^. To determine if JNK and ASK1 foci demonstrated liquid-like properties, we performed live-cell tracking and observed CRY-JNK2 and CRY-ASK1 were highly dynamic with neighbouring clusters undergoing fusion upon contact (Fig. 4A). Light induction of CRY-JNK2 and CRY-ASK1 clusters was prevented with 1,6-hexanediol, (Fig. 4, B and C) an aliphathic alcohol that disrupts protein condensates in mammalian cells ^32^. In contrast, Cry2_clust_ clusters were not perturbed by 1,6-hexanediol (Fig. 4D) consistent with their formation of stable homo-oligomers ^28^. In addition to increased numbers, CRY-JNK2 transitioned from elongated to spherical clusters with increased pHi, suggesting a pH-dependent change in their biophysical properties (Fig. 4E). Cytoplasmic CRY-JNK2 clusters at pHi 7.5 rapidly recovered 90% of maximum fluorescence intensity following photobleaching (Fig. 4, E and F). In comparison, CRY-ASK1 clusters recovered 40% of maximum fluorescence following photobleaching (Fig. 4, E to G). This indicates that JNK2 and a proportion of ASK1 within clusters are highly dynamic and rapidly exchange with the cytosol. In contrast, Cry2_clust_ clusters, regardless of their localization in cytoplasm or nucleus, and CRY-JNK2 at pHi 7.1 did not recover fluorescence following photobleaching (Fig. 4, E to G). Thus, under alkaline pHi conditions, JNK2 and ASK1 appear to be assembled as protein condensates.

**Fig. 4.**
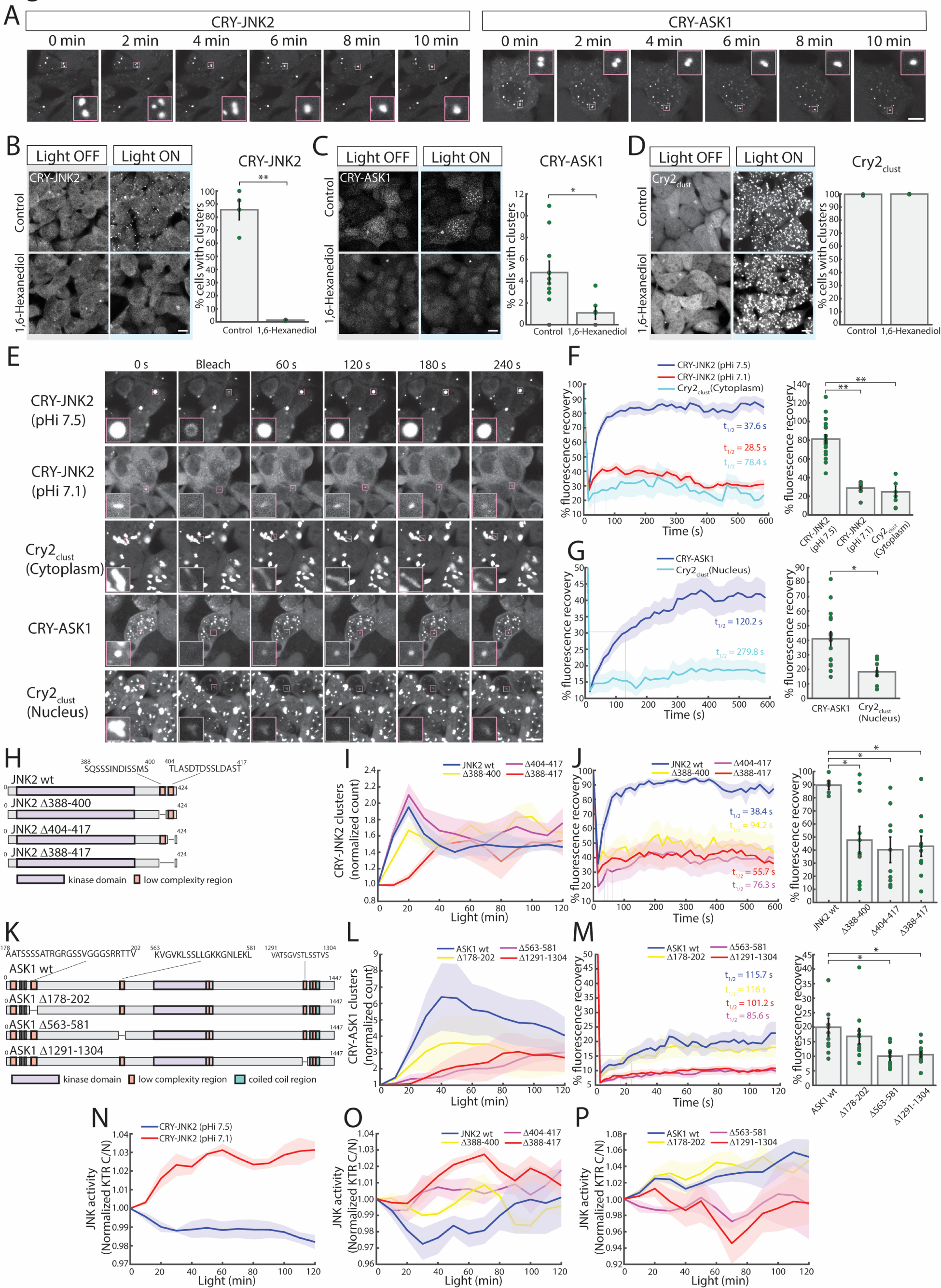
pHi regulates assembly of JNK2 and ASK1 into protein condensates. (**A**) Live imaging of HEK293 cells expressing CRY-JNK2 or CRY-ASK1 tracks fusion of distinct protein clusters over time. **(B)** HEK293 cells expressing CRY-JNK2 (n ≥ 3, per condition), (**C**) CRY-ASK1 (n ≥ 6, per condition) or (**D**) Cry2_clust_ (n ≥ 5, per condition) were pretreated with 1,6-Hexanediol (1%, 20 min) or a vehicle control prior to blue-light stimulation (30 min). (**E**) CRY-JNK2 clusters in HEK293 cells at pHi 7.1 or pHi 7.5 or CRY-ASK1 at pHi 7.5 were photobleached (5 s) and fluorescence recovery monitored for indicated time with fluorescence recovery after photobleached Cry2_clust_ assemblies (pHi 7.5) in the nucleus or cytoplasm for comparison. (**F**) Percentage fluorescence recovery of cytoplasmic Cry2_clust_ clusters (n = 7) or CRY-JNK2 clusters at pHi 7.1 (n = 10) or pHi 7.5 (n = 18). (**G**) Percentage fluorescence recovery of CRY-ASK1 (n = 15) or nuclear Cry2_clust_ (n = 7). (**H**) Schematic of JNK2 domains and deletion of identified low complexity regions. (**I**) Light induced clustering of CRY-JNK2 wild type (wt) and deletion mutants with removed low complexity regions (n = 3, per condition). (**J**) Fluorescence recovery of photobleached clusters of CRY-JNK2 and deletion mutants (n ≥5, per condition). (**K**) Schematic of ASK1 domains and deletion mutants with removed low complexity regions. (**L**) Light-induced clustering of CRY-ASK1 and deletion mutants with removed low complexity regions (n = 3, per condition). (**M**) Fluorescence recovery of photobleached clusters of CRY-ASK1 and deletion mutants (n ≥ 8, per condition). (**N**) Light-stimulated JNK activity (KTR) in HEK293 cells expressing CRY-JNK2 with pHi 7.1 or pHi 7.5 (n ≥ 10, per condition). (**O**) Light-stimulated JNK activity (KTR) in HEK293 cells expressing CRY-JNK2 and deletion mutants (n = 3, per condition). (**P**) Light-stimulated JNK activity (KTR) in HEK293 cells expressing CRY-ASK1 and deletion mutants (n = 3, per condition). Values represent mean±SEM. Percentage of maximal fluorescence recovery and half-time to maximal recovery (t_1/2_) indicated in **F**, **G**, **J** and **K**. Unpaired t test, *p < 0.05, **p < 0.001. Scale bars, 10 µm.

The formation of protein condensates is mediated by weak, multivalent interactions, commonly requiring low complexity or intrinsically disordered regions ^25,32^. SMART sequence analysis ^33^ identified regions of low complexity (aa 388-400 and 404-417) within the intrinsically disordered C-terminal tail of JNK2 (Fig. 4H). The deletion of JNK2 aa 388-400, but not aa 404-417, reduced the light-induced formation of CRY-JNK2 clusters (Fig. 4I) while the single deletion of either region was sufficient to blunt JNK2 fluorescence recovery following photobleach studies (Fig. 4J). SMART analysis identified a number of low complexity motifs within the ASK1 sequence and we selected three motifs (aa 178-202, 563-581 and 1291-1304) that were not located in structured regions for deletion (Fig. 4K). Deletions of each region individually was sufficient to reduce optogenetic stimulation of ASK1 clusters (Fig. 4L). However, fluorescence recovery of ASK1 clusters were specifically attenuated by Δ563-581 or Δ1291-1304 but not Δ178-202 (Fig. 4M). We conclude that these identified low complexity regions are required for the formation of JNK2 and ASK1 condensates at permissive pHi. Moreover, blue-light stimulation of Cry2_clust_-JNK2 triggered an increase in JNK activity when pHi was equilibrated to 7.1 but not at when pHi was increased to 7.5 (Fig. 4N). Light-stimulated activation of JNK at pHi 7.5 was marginally improved with deletion of either Δ388-400 or Δ404-417 low-complexity motifs (Fig. 4O) and while C-terminal JNK2 truncation removing both motifs resulted in light-stimulated JNK activation similar to that observed at more neutral permissive pHi (Fig. 4, N to O). Conversely, deletion of Δ563-581 or Δ1291-1304 but not Δ178-202 in ASK1 reduced light-stimulated JNK activation (Fig. 4P) coinciding with reduced fluorescence recovery of ASK1 clusters (Fig. 4M). Thus, ASK1 low-complexity regions facilitate condensate formation at increased pHi to promote activation of JNK while cytosolic alkalinization promotes JNK2 condensates that are antagonistic to signalling.

### Modelling of pH-regulated ASK1 and JNK2 predicts stress-context specific signal outputs

Our results suggest that the contrasting relationship between pHi and JNK activity with different stimuli may be explained by differential stress-activation of ASK1 and opposing effects of pHi on activities of kinases within the tiered JNK cascade. We examine this hypothesis, we employed a mathematical modelling approach to model the signal output from the pathway, incorporating pHi-regulated formation of JNK and ASK1 condensates and the differential ASK1 input caused by various stimuli (Fig. 5A). Our model builds upon the established tiered regulation of JNK by upstream MAP3Ks and negative feedback by DUSP1 phosphatases ^34^, which we further validated in our system using DUSP1 inhibitors, confirming that these resulted in enhanced stress-activation of JNK (Fig. S4A). We then incorporated in the model the pHi-assembled condensates that promote or attenuate the activity of ASK1 and JNK respectively, and differential input by ASK1 in response to stimuli as indicated in our studies (Fig. 2, 3 and Fig. S2). Our model also considered pHi as a dynamic signalling component that is increased by ASK1 and attenuated by JNK, based on the observed fluctuations of pHi in response to sorbitol and TNFα respectively (Fig. 1).

**Fig. 5.**
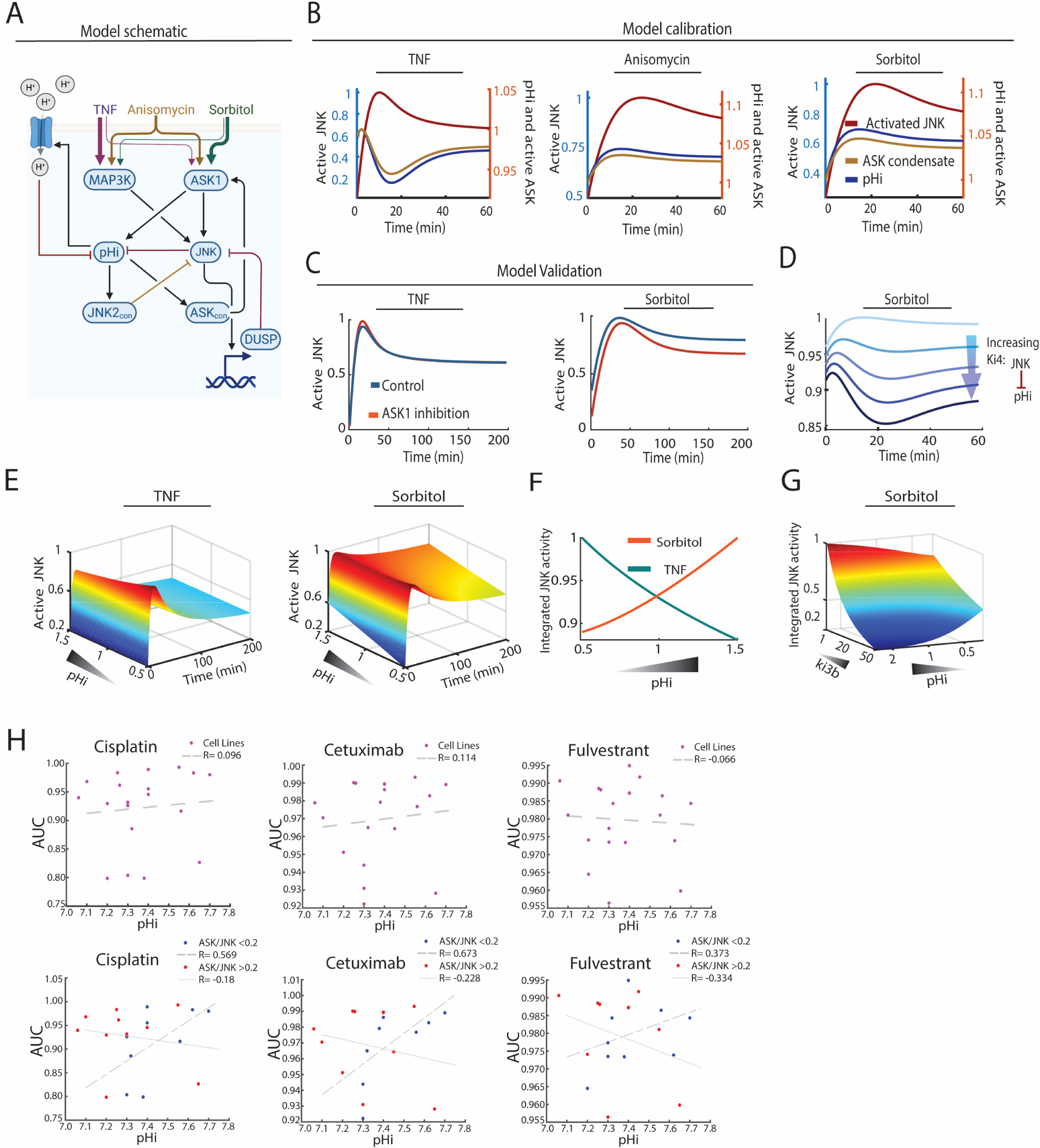
JNK cascade condensates and pHi predict signal output and functional consequences of the pathway in response to cell stress. (**A**) Model schematic illustrating the JNK MAPK pathway and its regulation by pHi in response to stimulation by stress stimuli. (**B**) Model simulations using the best-fitted parameter set show pHi and active JNK response dynamics following stress stimulation. TNFα-simulated response dynamics of pHi displayed an opposite trend to that of active JNK, while the trend in response to anisomycin or sorbitol was similar. (**C**) Independent model validation through simulation of active JNK in response to TNFα or hyperosmolarity under control conditions (blue) and with ASK1 inhibition (red). (**D**) The dynamic behaviour of pHi is controlled by increasing the inhibition rate of JNK on pHi (Ki4). This leads to the transition of pHi dynamics from upregulation to suppression, as observed after TNFα stimulation. (**E** to **F**) 3D time-dependent model simulations illustrate the temporal changes and opposite trends in JNK activity following TNFα and hyperosmotic stimulation at varying pHi levels. (**G**) JNK activity levels are dependent on pHi and *Ki3b* parameter (JNK inhibition rate by JNK2con). At low *Ki3b* values, there is a positive correlation between pHi levels and JNK activity under sorbitol stimulation. However, as *Ki3b* increases, the correlation between pHi levels and JNK activity becomes negative. (**H**) Correlation between pHi and cancer cell line sensitivity to anti-cancer drugs (cisplatin, cetuximab and fulvestrant) from GDSC database. The ratio of ASK1 to JNK2 in cancer cell types (blue dots ASK:JNK <0.2 and red dot indicate cells with ASK:JNK >0.2) delineated correlations between pHi and drug sensitivity. ‘R’ indicates Pearson correlation coefficient.

The model, depicted in Figure 5A, was formulated using ordinary differential equations (ODEs) and implemented in MATLAB. This effectively characterises the network interactions as a series of ODEs based on kinetic laws (see Methods for a detailed model description). We calibrated the model using the experimental data of JNK and pHi dynamics in response to TNFα, anisomycin and sorbitol (Fig. 5B). Simulation of stress treatments with the calibrated model recapitulated the experimentally observed changes in JNK activity, pHi and formation of ASK1 condensates (Fig. 5C and Fig. 1 and 2, Fig. S1). By validating our model using independent data, we found that it accurately predicted the effect of ASK1 inhibition on JNK activity stimulated by sorbitol but not TNFα (Fig. 5B). Using the model, we explored the mechanism regulating pHi dynamics in response to different stress stimuli. For this, we undertook a sensitivity analysis by systematically perturbing model parameters (see Methods) and simulating the effect on pHi dynamics in response to stress treatment. The results identified the strength of JNK-induced pHi acidification (*Ki4*) as the primary determinant of varying pHi dynamics after stress treatment. At low *Ki4,* pHi is promoted by sorbitol treatment, but a higher *Ki4* level switches this regulation from promotion to suppression (Fig. 5D).

Our model also accurately predicted the relationship between pHi and JNK activation the strength under different stress types (Fig. 5, E and F). Notably, simulations showed a negative correlation between pHi and JNK activity under TNFα treatment, but a positive correlation under sorbitol treatment. Next, we conducted a model sensitivity analysis to decipher the regulatory mechanisms affecting the pHi-JNK activity relationship (Fig. 5G and Fig. S4, B and C). The results indicate that the strength of JNK inhibition by JNK2con (*ki3b*) under sorbitol treatment, and the activation rate of ASK by ASKcon (*kf1d*) under TNFα stimulation are the key parameters in governing the pHi-JNK activity relationship (Fig. S4, B and C). For example, Figure 5G depicts a shift from a positive correlation between pHi and JNK activity to a negative association, induced by increasing the *ki3b* parameter values (see Fig. S4D for a similar analysis for *kf1d*). Together, our findings demonstrate how the dynamic interplay between ASK1 and JNK activities, alongside pHi-regulated kinase condensate formation, can explain the varied impact of pHi on stress-induced JNK pathway signalling.

We next hypothesized that knowledge of pHi and the kinases involved in the JNK network could help predict the strength of the pathway’s signal output, which determines cellular outcomes. To test this this idea, we turned to an analysis of cancer cell sensitivity to anti-tumour drugs in the GDSC database. Specifically, drugs like cisplatin, cetuximab, and fulvestrant, known to exert anti-tumour effects through JNK pathway activation, were considered ^35,36^. Although initial observation showed no clear correlation between drug sensitivity and pHi, a nuanced analysis revelaed reduced drug sensitivity at higher pHi among cancer cell types with low ASK1 expression relative to JNK2 (Fig. 5H). This is consistent with weak JNK signal outputs that are not conducive to programmed death induction. In contrast, cancer cells with high ASK1 levels relative to JNK2 showed a modest positive pHi correlation with drug sensitivity (Fig. 5H), suggesting that in this subgroup, elevated pHi enhances JNK activation leading to cell death. Thus, our findings indicate that pHi can help resolve the contextual function of JNK signals in cancer cell responses to anti-cancer therapy and other pathophysiological contexts where the regulatory roles of JNK have remained obscure.

## Discussion

The transduction of biochemical signals in cells relies on ions, nucleotides and phopsholipids^37^. Our work indicates that cytosolic protons (H^+^) are intimately involved in signal transduction and should be considered as an integral component of intracellular regulatory networks. While pHi is tightly regulated to guard against dramatic fluctuations that can damage proteins and trigger cell death, it is increasingly evident that localized physiological pHi changes have specific cell regulatory effects ^12–14^. Exocytosed protons participate in intercellular communication, including direct negative regulation of calcium channels during synaptic transmission ^38^, or positive signal feedback through stimulation of acid-sensing ion channels ^39^. In yeast, intracellular protons have been shown to act as second messengers in the glucose activation of protein kinase A ^9^ and are coupled with GPCR activation for coincidence signal detection ^40^. Here, we demonstrated in mammalian cells that pHi changes are integrated within signal networks to determine the strength of JNK activation in response to specific stimuli. The JNK pathway has diverse, highly contextual functions that is dependent on how JNK signals are interpreted by cells ^2^. A pHi-regulated contextual JNK response may, therefore, serve to provide precise outcomes that are dependent on how signal inputs are generated.

We find that pHi alters signal transduction by modifying the biophysical properties and activity of kinases embedded within the JNK pathway. Our optogenetic seeding of ASK1 and JNK2 condensates revealed that the relative alkalinization of the cytosolic environment promoted protein interactions. This is reminiscent of pHi-dependent conformational changes and oligomerization of Bcl-xL and Bax apoptotic proteins ^41,42^, mediated through altered protonation states of charged amino acids. Histidine residues are typically implicated as pH sensors due to the near physiological pKa (∼6.0) of their imidazole side group ^10^. However, the pKa of other ionizable residues are known to shift dramatically towards physiological pH, and this is partly dependent on their position and topology within protein structures ^43^. Along these lines, the pH-dependent phase transition of a yeast prion protein, Sup35, is regulated by protonation of a linear cluster of glutamic acid residues ^44^. In this study, we identified regions in the C-terminal tail of JNK2 that were enriched with polar serine and negatively charged aspartate residues and were required for pH-dependent condensate formation. Similarly, ASK1 low complexity domains involved in phase separation were comprised of positively charged lysines. This suggests that these domains may respond to increased pHi to promote kinase interactions triggering condensate formation. It remains a possibility that evolutionarily conserved histidines in JNK2 and ASK1 may further contribute to the observed kinase responses to pHi. Notably, the mechanisms that underlie pH-dependent kinase responses may lead to new approaches to target kinase activity and modulate signal transduction in cells.

In response to stress stimuli, pHi is known to regulate the phase transition of RNA-binding G3BP1, DEAD-Box helicase and poly(A)-binding proteins ^45–47^. However, the pH-dependent phase transitions of these proteins occur at pHi<6.0 which is much lower than physiological pH. Under such conditions, phase separation represents an adaptive mechanism to sequester disrupted proteins into phase separated granules to maintain protein function and cell survival^48^. In contrast, JNK2 and ASK1 undertook phase transitions at relatively alkaline but physiological pHi between 7.5 and 8.0 which indicate mild changes in pHi function as signal regulators. In optogenetic studies, while light-clustering of JNK2 at relatively acidic pHi increased activity, the liquid-like phase transition at alkaline pHi inhibited signalling. So we speculate that JNK2 condensates are not conducive to trans-autoactivation ^26^ or are spatially separated from their downstream targets including nuclear transcription factors ^49^. The phase transition of ASK1 has the opposing effect of enhancing downstream signalling to JNK. Thus, condensate formation may facilitate a quaternary state of ASK1 that enhances kinase activation ^50^ or concentrate ASK1 with its targets perhaps by bringing together ASK1-binding signal scaffolds ^51^. This is consistent with phase-separated condensates acting as subcellular platforms that organize kinase signalling ^31^. Thus, opposing effects of phase separation on the activity of protein kinases organized within tiered cascades create a complex signal feedback system that defines signal output. Our mathematical modelling of the JNK pathway, which accounts for stimuli-induced alterations in pHi and consequent condensate formation of ASK1 and JNK, was able to recapitulate experimental findings related to changes in JNK activity, pHi, and ASK1 condensate formation. Furthermore, it could predict JNK signal outputs at specific pHi under various stress conditions and identify potential regulatory interactions controlling the pHi-JNK correlation. Our model provides a robust tool that will aid in the future delineation of the complex and diverse functions of JNK and related tiered kinase pathways under different conditions.

Our studies indicate that the pHi-JNK relationship may have benefit in delineating signal regulation in cancer and predicting responses to anti-cancer therapy. JNK signals promote tumour growth and metastasis and mediates cell death in response to anti-cancer drugs ^4^. Precise measures of pHi may delineate the role of JNK signalling in tumours, and an understanding of the positive or negative association between JNK signal capacity and pHi could be exploited therapeutically to prevent oncogenic signals while promoting JNK-killing of cancer cells in response to chemo- or radiotherapy. More broadly, the hallmark inversion of the pH gradient in cancer cells suggest that inhibitors of proton pumps and exchangers may improve chemosensitivity of tumours ^52^. However, trials have returned mixed results with adverse patient outcomes reported when proton pump inhibitors were combined with kinase-targeting drugs or immune-checkpoint inhibitors ^53,54^. While this is in part due to drug-drug interactions ^55^, an improved understanding of the association between increased pHi and kinase signal regulation of cancer progression appears to be necessary to optimize pHi targeting as a therapeutic strategy. In addition to cancer, dysregulation of JNK signalling is linked to neurodegeneration, cardiac remodeling, metabolic and inflammatory disorders ^1^, where our findings may help decipher the contextual consequences of the pathway.

## Methods

### Literature and database analysis

Dataset for intracellular pHi measurements for Fig.5H and Fig. S1A was generated from the previously published data and presented in the Table S1. Data for the efficiency of anticancer drugs (AS601245, JNK inhibitor VIII, JNK 9L, cisplatin, cetuximab, and fulvestrant) as AUC of viability plot were obtained from Genomics of Drug Sensitivity in Cancer (GDSC) database^56^ (www.cancerRxgene.org). Expression levels of JNK2 and ASK1 in a panel of cell lines were obtained from the Human Protein Atlas^57^ (www.proteinatlas.org).

### DNA Constructs

Plasmids were constructed using standard restriction enzymes cloning. pPBbsr-JNKKTR-mCherry plasmid containing JNK activity reporter (JNK-KTR) was obtained from Addgene (#115493). mCherry in this plasmid was exchanged with iRFP713 (from Addgene #111510) fluorophore using NotI and SalI enzymes. ClopHensor plasmid was obtained from Addgene (#25938). JNK2α2 and ASK1 were amplified from pCDNA3 Flag Jnk2a2 (Addgene #13755) and pCMV6-Myc-MAP3K5 (ASK1) (Origen #RC209913). PCR amplified constructs were inserted into mCherry-CRY2clust (Addgene #105624), using BsrGI/HpaI sites to obtain mCherry tagged JNK2α2 and SgrAI/HpaI sites to obtain mCherry tagged ASK1 constructs. Amplified JNK2α2 and ASK1 were also inserted into mCherry-CRY2clust using BsrGI/BspEI sites and SmaI/HpaI sites respectively to obtain CRY-JNK2α2 and CRY-ASK1. Obtained constructs containing CMV promoter, mCherry, JNK2α2 or ASK1, and Cry2_clust_ were then cloned into PiggyBac backbone (a gift from M. Jones) using SpeI/PmeI sites. Low complexity regions of JNK2α2 and ASK1 were deleted using Q5 Site-Directed Mutagenesis (NEB). Oligonucleotides used for cloning and site-directed mutagenesis are listed in the Table S2.

### Cell Maintenance, transfection, and stable cell line generation

U87MG, MCF7, HEK293, SH-SY5Y, and U2OS cell lines were purchased form American Type Culture Collection (ATCC). U87MG, MCF7, HEK293, U2OS were cultured in DMEM (Gibco) supplemented with 10% FBS (Gibco) and 1% penicillin/streptomycin (Gibco) at 37°C with 5% CO2 in a humidified incubator. SH-SY5Y was cultured in the same condition but with 15% FBS. For transfections cells were seeded at the 8-well chamber slides (ibidi) at the 80% confluency and next day transfected with 0.5 µg plasmid and Lipofectamine (Invitrogen) at the ratio of 1:6. Cells were used for imaging 24 hours after transfection. For stable cell lines generation, cells were seeded at the 12-well plate and transfected with plasmids (0.5 µg each) using Lipofectamine. For generation of cells lines with CRY-JNK2α2 and CRY-ASK1, cells were transfected with plasmids of interests and transposases (with the transposon to transposase ratio of 1:3) using Lipofectamine. 3 days after transfection cell were treated with antibiotics according to resistance, 1 mg/ml G418 (Sigma-Aldrich) and/or 5 µg/ml blasticidin (Sigma-Aldrich). After 7 days of selection, cells were plated at the 96 well plates with a density of 1 cell per well. Obtained monoclonal cell populations with targeted proteins were cultured for 2 weeks prior using in the experiments. The list of generated cell lines for this study are presented at Table S3.

### Stress treatment

8-well chamber slides were coated with fibronectin in PBS (2.5 µg/cm^2^, Sigma-Aldrich) for 1 hour. Cells were seeded at the coated chamber slides with a density 200000 cells per well and the next day four hours before imaging cells were stained with 1 µg/ml Hoechst 33342 for 30 min at 37°C and washed 3 times for 10 min each with Live Cell Imaging Solution (Invitrogen). When applicable, cells were pre-treated with inhibitors (10 µM ASK inhibitor or 1 µM JNK inhibitor VIII or 10 µM BCI (Sigma-Aldrich)) and then treated with 10 ng/ml TNFα (Sigma-Aldrich), 10 ng/ml anisomycin (Sigma-Aldrich) or 150 mM sorbitol (Sigma-Aldrich). For modulation of intracellular pH, before imaging cells were treated with 50 µM Bafilomycin A1 (Sigma-Aldrich) or 50 mM NH_4_Cl (Supelco) or 10 µM nigericin (Invitrogen). Nigericin was added to cells by exchanging cell media with a new one with pH 7.0 and containing nigericin. For the duration of all imaging experiments cells were kept at 37 °C and 5% CO2. Images of JNK-KTR translocation were obtained with 633 nm laser, Leica DMi8 SP8. JNK activity was calculated as the ratio between intensities of fluorescence of JNK-KTR in the cytoplasm and in the nucleus (C/N ratio). Values of ratios were normalized by the value of C/N ratio of JNK-KTR in cells before stress treatment was applied. Intensities and C/N ratio of JNK-KTR were measured using CellProfiler 4.0.7, data was than normalized and plotted in MATLAB R2021a.

### Intracellular pH sensing

Intracellular pH in U87MG, MCF7, HEK293, SH-SY5Y, and U2OS cell lines was measured using chemical dye pHrodo-Green AM (Invitrogen) and genetically encoded pH and chloride indicator ClopHensor. To measure intracellular pH with pHrodo, U87MG, MCF7, HEK293, SH-SY5Y, U2OS cells were seeded at the 8-well chamber slides with a density 200000 cells per well and stained with pHrodo according to the manufacturer’s standard protocol (Invitrogen). At least four hours after staining cells were imaged using a 488 nm laser, Leica DMi8 SP8. Intensity of pHrodo was used to calculate pH. Intracellular pH was also measured using the genetically encoded pH and chloride sensor ClopHensor. ClopHensor was transiently expressed in U87MG, MCF7, SH-SY5Y, and U2OS, and stably expressed in HEK293 cells. ClopHensor fluorescence was imaged with 458 nm (cyan emission) and 488 nm (green emission) laser, Leica DMi8 SP8. The ratio between fluorescent intensities at 488 nm and 458 nm was used to calculate pH. For pH quantification calibration curves were built for all cell lines for pHrodo staining, and for ClopHensor expressed in HEK293 cells. To build a calibration curves, intracellular pH calibration buffer kit (Invitrogen) buffers with pH 4.5, 5.5, 6.5, and 7.5 and additional pH buffers made by adjustments of pH with 1 mM NaOH (ChemSupply Australia) were used. pH for pHrodo stained cells was defined using linear regression analysis (MATLAB R2021a). The value for pHi for ClopHensor was calculated according to equation: 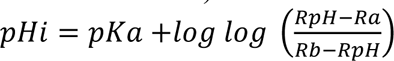, where pKa = 6.75, Ra = 5.8, Rb = 12, RpH is a green-to-cyan ratio. Intensities of pHrodo and cyan-to-green ratio of ClopHensor were measured using CellProfiler 4.0.7, data was than plotted in MATLAB R2021a.

### Optogenetic modulation of JNK2α2 and ASK1 clustering

HEK293 cells stably expressing CRY-JNK2, CRY-ASK1, and Cry2_clust_ wear seeded at the 8-well chamber slides with a density 200000 cells per well (if other not specified). Mutated variants of CRY-JNK2 and CRY-ASK1 and controls were transiently expressed in HEK293 cells. To induce clustering of CRY-JNK2, CRY-ASK1, and Cry2_clust_, cells were illuminated with blue light (470 nm, 21 mW, Mightex). When applicable, prior photoactivation cells were treated with 50 µM Bafilomycin A1 (Sigma-Aldrich), 50 mM NH_4_Cl (Supelco), 10 µM nigericin or 1,6-hexanediol (Sigma-Aldrich) with a final concentration of 1% (v/v). Images of CRY-JNK2, CRY-ASK1, and Cry2_clust_ clusters were acquired with 561 nm laser, Leica DMi8 SP8. The number of clusters per cell were measured using CellProfiler 4.0.7, data was than normalized (to the number of clusters before illumination with blue light) and plotted in MATLAB R2021a.

### FRAP

FRAP assays were performed using Leica DMi8 SP8. Clusters were bleached with 80% intensity of 561 nm laser for 5 s, with the following image acquiring every 15 sec for 10 min. Intensities of fluorescence was extracted using ImageJ and fitted using MATLAB R2021a. The half-time of recovery (t_1/2_) was calculated as the time required to recover half maximum recovery intensity of fluorescence. Half of maximum recovered fluorescence intensity was calculated as: *I_1/2_ = (I_sat_ + I_0_)/2*, where *I_sat_* is an intensity of bleached area after recovery at the maximum saturation point, and *I_0_* is an intensity of bleached area immediately after bleaching.

### Statistical analysis

All statistical analyses were carried out with MATLAB R2021a. Data were analyzed using unpaired t test. P-values < 0.05 were considered statistically discernible (*p < 0.05; **p < 0.001).

### Mathematical Modelling

In this study, we developed a mechanistic model that reflects the activity of the JNK pathway dependent on pHi. This model was developed using ordinary differential equations (ODEs), a type of equation used in chemical kinetics. The full set of reaction rates can be found in Table S4. In Supplementary Information 1, we provide the best-fitted parameter sets used for the simulations, which were derived from the model fitting (calibration) process. Model implementation and simulation were performed using MATLAB (The MathWorks. Inc. 2020b) and the IQM toolbox (http://www.intiquan.com/intiquan-tools/). Model fitting was done by minimising the objective function J below, which quantifies the discrepancies between the fitted experimental data and corresponding model simulations:

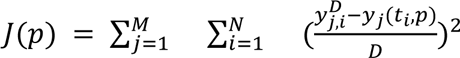

Here, M denotes the number of experimental data sets used for fitting; N is the number of time points within each set; *y_j_(t_i_*, *p)* indicates simulation value of component j at the time point *t_i_* with parameter set *p* and 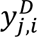 is the experimental data of component *j* at time point *t_i_*. D also denotes the standard deviation of the experimental data.

Figures 1B, 1G and Supplementary Figure 1G were utilized to calibrate the model based on experimental data. The objective function j was minimized by estimating the parameter values. The parameter estimation process was carried out using the Global Optimization Toolbox in MATLAB, specifically employing the ga function. A population size of 400 and a generation number of 800 were used during this parameter estimation process. The code and parameter sets have been stored in Github (https://github.com/NguyenLab-IntegratedNetworkModeling/JNK_pHi).

### Sensitivity analysis

Model-based sensitivity analysis was employed to investigate the mechanism governing the correlation between pHi and JNK activity. To do this, we devised a metric that quantifies the relationship between pHi and JNK activity, defined as the slope of a simulated pHi-JNK dose-response curve. This curve was generated by varying the pHi within a broad range (0.5 to 1.5), and the resulting JNK activity was determined using our model (as shown in Fig. 5F). This slope metric was then applied in the sensitivity analysis. Here, each kinetic parameter in the model was systematically altered within an extensive range (from 0.001 to 1000 times its nominal value), and the effect on the slope metric was calculated. Finally, the parameters were ranked from those with the most significant impact to those with the least (as seen fig S4, B and C).

## Acknowledgments

We would like to thank School of Biomedical Science Imaging and Analytical facility and Dr Shaun Walters for microscopy assistance. We thank Dr Sebastian Furness, Prof Walter Thomas, A/Prof Sean Millard and Dr Uda Ho for helpful discussion and for providing feedback on the manuscript.

## Author contributions

Conceptualization: YD, DCHN Methodology: YD, MG Investigation: YD, DCHN Visualization: YD, MG Funding acquisition: DCHN Supervision: DCHG

Project administration: DCHN, LKN Writing – original draft: DCHG

Writing – review & editing: YD, MG, LKN

## Data and materials availability

All data are available in the main text or the supplementary materials, files and from the authors upon request.

## References

1 Bogoyevitch, M. A., Ngoei, K. R., Zhao, T. T., Yeap, Y. Y. & Ng, D. C. c-Jun N-terminal kinase (JNK) signaling: recent advances and challenges. Biochim Biophys Acta 1804, 463–475 (2010). 10.1016/j.bbapap.2009.11.002

2 Davis, R. J. Signal transduction by the JNK group of MAP kinases. Cell 103, 239–252 (2000). 10.1016/s0092-8674(00)00116-1

3 La Marca, J. E. & Richardson, H. E. Two-Faced: Roles of JNK Signalling During Tumourigenesis in the Drosophila Model. Front Cell Dev Biol 8, 42 (2020). 10.3389/fcell.2020.00042

4 Tournier, C. The 2 Faces of JNK Signaling in Cancer. Genes Cancer 4, 397–400 (2013). 10.1177/1947601913486349

5 Ventura, J. J. et al. Chemical genetic analysis of the time course of signal transduction by JNK. Mol Cell 21, 701–710 (2006). 10.1016/j.molcel.2006.01.018

6 Brancho, D. et al. Role of MLK3 in the regulation of mitogen-activated protein kinase signaling cascades. Mol Cell Biol 25, 3670–3681 (2005). 10.1128/MCB.25.9.3670-3681.2005

7 Saitoh, M. et al. Mammalian thioredoxin is a direct inhibitor of apoptosis signal-regulating kinase (ASK) 1. EMBO J 17, 2596–2606 (1998). 10.1093/emboj/17.9.2596

8 Watanabe, K., Umeda, T., Niwa, K., Naguro, I. & Ichijo, H. A PP6-ASK3 Module Coordinates the Bidirectional Cell Volume Regulation under Osmotic Stress. Cell Rep 22, 2809–2817 (2018). 10.1016/j.celrep.2018.02.045

9 Dechant, R. et al. Cytosolic pH is a second messenger for glucose and regulates the PKA pathway through V-ATPase. EMBO J 29, 2515–2526 (2010). 10.1038/emboj.2010.138

10 White, K. A. et al. beta-Catenin is a pH sensor with decreased stability at higher intracellular pH. J Cell Biol 217, 3965–3976 (2018). 10.1083/jcb.201712041

11 Roos, A. & Boron, W. F. Intracellular pH. Physiol Rev 61, 296–434 (1981). 10.1152/physrev.1981.61.2.296

12 Spear, J. S. & White, K. A. Single-cell intracellular pH dynamics regulate the cell cycle by timing G1 exit and the G2 transition. J Cell Sci (2023). 10.1242/jcs.260458

13 Srivastava, J. et al. Structural model and functional significance of pH-dependent talin-actin binding for focal adhesion remodeling. Proc Natl Acad Sci U S A 105, 14436–14441 (2008). 10.1073/pnas.0805163105

14 Ulmschneider, B. et al. Increased intracellular pH is necessary for adult epithelial and embryonic stem cell differentiation. J Cell Biol 215, 345–355 (2016). 10.1083/jcb.201606042

15 Webb, B. A., Chimenti, M., Jacobson, M. P. & Barber, D. L. Dysregulated pH: a perfect storm for cancer progression. Nat Rev Cancer 11, 671–677 (2011). 10.1038/nrc3110

16 Isom, D. G. et al. Protons as second messenger regulators of G protein signaling. Mol Cell 51, 531–538 (2013). 10.1016/j.molcel.2013.07.012

17 Li, X., Augustine, A., Sun, D., Li, L. & Fliegel, L. Activation of the Na(+)/H(+) exchanger in isolated cardiomyocytes through beta-Raf dependent pathways. Role of Thr(653) of the cytosolic tail. J Mol Cell Cardiol 99, 65–75 (2016). 10.1016/j.yjmcc.2016.08.014

18 Verkest, C., Diochot, S., Lingueglia, E. & Baron, A. C-Jun N-Terminal Kinase Post-Translational Regulation of Pain-Related Acid-Sensing Ion Channels 1b and 3. J Neurosci 41, 8673–8685 (2021). 10.1523/JNEUROSCI.0570-21.2021

19 Miura, H., Kondo, Y., Matsuda, M. & Aoki, K. Cell-to-cell heterogeneity in p38-mediated cross-inhibition of JNK causes stochastic cell death. Cell reports 24, 2658–2668 (2018).

20 Regot, S., Hughey, J. J., Bajar, B. T., Carrasco, S. & Covert, M. W. High-sensitivity measurements of multiple kinase activities in live single cells. Cell 157, 1724–1734 (2014).

21 Reinhard, C., Shamoon, B., Shyamala, V. & Williams, L. T. Tumor necrosis factor alpha-induced activation of c-jun N-terminal kinase is mediated by TRAF2. EMBO J 16, 1080–1092 (1997). 10.1093/emboj/16.5.1080

22 Galcheva-Gargova, Z., Derijard, B., Wu, I. H. & Davis, R. J. An osmosensing signal transduction pathway in mammalian cells. Science 265, 806–808 (1994). 10.1126/science.8047888

23 Yoshimori, T., Yamamoto, A., Moriyama, Y., Futai, M. & Tashiro, Y. Bafilomycin A1, a specific inhibitor of vacuolar-type H(+)-ATPase, inhibits acidification and protein degradation in lysosomes of cultured cells. J Biol Chem 266, 17707–17712 (1991).

24 Huc, L. et al. c-Jun NH2-terminal kinase-related Na+/H+ exchanger isoform 1 activation controls hexokinase II expression in benzo(a)pyrene-induced apoptosis. Cancer Res 67, 1696–1705 (2007). 10.1158/0008-5472.CAN-06-2327

25 Jin, X. et al. Effects of pH alterations on stress- and aging-induced protein phase separation. Cell Mol Life Sci 79, 380 (2022). 10.1007/s00018-022-04393-0

26 Nitta, R. T., Chu, A. H. & Wong, A. J. Constitutive activity of JNK2 alpha2 is dependent on a unique mechanism of MAPK activation. J Biol Chem 283, 34935–34945 (2008). 10.1074/jbc.M804970200

27 Trevelyan, S. J. et al. Structure-based mechanism of preferential complex formation by apoptosis signal-regulating kinases. Sci Signal 13 (2020). 10.1126/scisignal.aay6318

28 Park, H. et al. Optogenetic protein clustering through fluorescent protein tagging and extension of CRY2. Nat Commun 8, 30 (2017). 10.1038/s41467-017-00060-2

29 Galkina, S. I., Sud’ina, G. F., Dergacheva, G. B. & Margolis, L. B. Regulation of intracellular pH by cell-cell adhesive interactions. FEBS Lett 374, 17–20 (1995). 10.1016/0014-5793(95)00969-g

30 Thomas, J. A., Buchsbaum, R. N., Zimniak, A. & Racker, E. Intracellular pH measurements in Ehrlich ascites tumor cells utilizing spectroscopic probes generated in situ. Biochemistry 18, 2210–2218 (1979). 10.1021/bi00578a012

31 Lopez-Palacios, T. P. & Andersen, J. L. Kinase regulation by liquid-liquid phase separation. Trends Cell Biol (2022). 10.1016/j.tcb.2022.11.009

32 Alberti, S., Gladfelter, A. & Mittag, T. Considerations and Challenges in Studying Liquid-Liquid Phase Separation and Biomolecular Condensates. Cell 176, 419–434 (2019). 10.1016/j.cell.2018.12.035

33 Schultz, J., Milpetz, F., Bork, P. & Ponting, C. P. SMART, a simple modular architecture research tool: identification of signaling domains. Proc Natl Acad Sci U S A 95, 5857–5864 (1998). 10.1073/pnas.95.11.5857

34 Tomida, T., Takekawa, M. & Saito, H. Oscillation of p38 activity controls efficient pro-inflammatory gene expression. Nat Commun 6, 8350 (2015). 10.1038/ncomms9350

35 Eke, I. et al. EGFR/JIP-4/JNK2 signaling attenuates cetuximab-mediated radiosensitization of squamous cell carcinoma cells. Cancer Res 73, 297–306 (2013). 10.1158/0008-5472.CAN-12-2021

36 Zanke, B. W. et al. The stress-activated protein kinase pathway mediates cell death following injury induced by cis-platinum, UV irradiation or heat. Curr Biol 6, 606–613 (1996). 10.1016/s0960-9822(02)00547-x

37 Newton, A. C., Bootman, M. D. & Scott, J. D. Second Messengers. Cold Spring Harb Perspect Biol 8 (2016). 10.1101/cshperspect.a005926

38 DeVries, S. H. Exocytosed protons feedback to suppress the Ca2+ current in mammalian cone photoreceptors. Neuron 32, 1107–1117 (2001). 10.1016/s0896-6273(01)00535-9

39 Waldmann, R., Champigny, G., Bassilana, F., Heurteaux, C. & Lazdunski, M. A proton-gated cation channel involved in acid-sensing. Nature 386, 173–177 (1997). 10.1038/386173a0

40 Kapolka, N. J. et al. Proton-gated coincidence detection is a common feature of GPCR signaling. Proc Natl Acad Sci U S A 118 (2021). 10.1073/pnas.2100171118

41 Bhat, V. et al. Acidic pH promotes oligomerization and membrane insertion of the BclXL apoptotic repressor. Arch Biochem Biophys 528, 32–44 (2012). 10.1016/j.abb.2012.08.009

42 Cartron, P. F., Oliver, L., Mayat, E., Meflah, K. & Vallette, F. M. Impact of pH on Bax alpha conformation, oligomerisation and mitochondrial integration. FEBS Lett 578, 41–46 (2004). 10.1016/j.febslet.2004.10.080

43 Isom, D. G. & Dohlman, H. G. Buried ionizable networks are an ancient hallmark of G protein-coupled receptor activation. Proc Natl Acad Sci U S A 112, 5702–5707 (2015). 10.1073/pnas.1417888112

44 Franzmann, T. M. et al. Phase separation of a yeast prion protein promotes cellular fitness. Science 359 (2018). 10.1126/science.aao5654

45 Guillen-Boixet, J. et al. RNA-Induced Conformational Switching and Clustering of G3BP Drive Stress Granule Assembly by Condensation. Cell 181, 346–361 e317 (2020). 10.1016/j.cell.2020.03.049

46 Iserman, C. et al. Condensation of Ded1p Promotes a Translational Switch from Housekeeping to Stress Protein Production. Cell 181, 818–831 e819 (2020). 10.1016/j.cell.2020.04.009

47 Riback, J. A. et al. Stress-Triggered Phase Separation Is an Adaptive, Evolutionarily Tuned Response. Cell 168, 1028–1040 e1019 (2017). 10.1016/j.cell.2017.02.027

48 Kroschwald, S. & Alberti, S. Gel or Die: Phase Separation as a Survival Strategy. Cel l168, 947–948 (2017). 10.1016/j.cell.2017.02.029

49 Maik-Rachline, G., Zehorai, E., Hanoch, T., Blenis, J. & Seger, R. The nuclear translocation of the kinases p38 and JNK promotes inflammation-induced cancer. Sci Signal 11 (2018). 10.1126/scisignal.aao3428

50 Liu, H., Nishitoh, H., Ichijo, H. & Kyriakis, J. M. Activation of apoptosis signal-regulating kinase 1 (ASK1) by tumor necrosis factor receptor-associated factor 2 requires prior dissociation of the ASK1 inhibitor thioredoxin. Mol Cell Biol 20, 2198–2208 (2000). 10.1128/MCB.20.6.2198-2208.2000

51 Matsuura, H. et al. Phosphorylation-dependent scaffolding role of JSAP1/JIP3 in the ASK1-JNK signaling pathway. A new mode of regulation of the MAP kinase cascade. J Biol Chem 277, 40703–40709 (2002). 10.1074/jbc.M202004200

52 Koltai, T. Targeting the pH Paradigm at the Bedside: A Practical Approach. Int J Mol Sci 21 (2020). 10.3390/ijms21239221

53 Chang, Y. et al. The Association between Baseline Proton Pump Inhibitors, Immune Checkpoint Inhibitors, and Chemotherapy: A Systematic Review with Network Meta-Analysis. Cancers (Basel) 15 (2022). 10.3390/cancers15010284

54 Sharma, M. et al. The concomitant use of tyrosine kinase inhibitors and proton pump inhibitors: Prevalence, predictors, and impact on survival and discontinuation of therapy in older adults with cancer. Cancer 125, 1155–1162 (2019). 10.1002/cncr.31917

55 van Leeuwen, R. W., van Gelder, T., Mathijssen, R. H. & Jansman, F. G. Drug-drug interactions with tyrosine-kinase inhibitors: a clinical perspective. Lancet Oncol 15, e315–326 (2014). 10.1016/S1470-2045(13)70579-5

56 Yang, W. et al. Genomics of Drug Sensitivity in Cancer (GDSC): a resource for therapeutic biomarker discovery in cancer cells. Nucleic acids research 41, D955–D961 (2012).

57 Uhlén, M. et al. Tissue-based map of the human proteome. Science 347, 1260419 (2015).

58 Belhoussine, R., Morjani, H., Sharonov, S., Ploton, D. & Manfait, M. Characterization of intracellular pH gradients in human multidrug-resistant tumor cells by means of scanning microspectrofluorometry and dual-emission-ratio probes. International journal of cancer 81, 81–89 (1999).

59 Hulikova, A., Vaughan-Jones, R. D. & Swietach, P. Dual role of CO2/HCO3(-) buffer in the regulation of intracellular pH of three-dimensional tumor growths. J Biol Chem 286, 13815–13826 (2011). 10.1074/jbc.M111.219899

60 Sennoune, S. R. et al. Vacuolar H+-ATPase in human breast cancer cells with distinct metastatic potential: distribution and functional activity. American journal of physiology-cell physiology 286, C1443–C1452 (2004).

61 Hulikova, A., Aveyard, N., Harris, A. L., Vaughan-Jones, R. D. & Swietach, P. Intracellular carbonic anhydrase activity sensitizes cancer cell pH signaling to dynamic changes in CO2 partial pressure. Journal of Biological Chemistry 289, 25418–25430 (2014).

62 Hanson, D. J. et al. Effective impairment of myeloma cells and their progenitors by blockade of monocarboxylate transportation. Oncotarget 6, 33568 (2015).

63 Baba, M., Inoue, M., Itoh, K. & Nishizawa, Y. Blocking CD147 induces cell death in cancer cells through impairment of glycolytic energy metabolism. Biochemical and biophysical research communications 374, 111–116 (2008).

64 Cianchi, F. et al. Selective inhibition of carbonic anhydrase IX decreases cell proliferation and induces ceramide-mediated apoptosis in human cancer cells. Journal of Pharmacology and Experimental Therapeutics 334, 710–719 (2010).

65 McLean, L. A., Roscoe, J., Jørgensen, N. K., Gorin, F. A. & Cala, P. M. Malignant gliomas display altered pH regulation by NHE1 compared with nontransformed astrocytes. American Journal of Physiology-Cell Physiology 278, C676–C688 (2000).

66 Ordway, B. et al. Targeting of evolutionarily acquired cancer cell phenotype by exploiting pHi-metabolic vulnerabilities. Cancers 13, 64 (2020).

67 Rich, I. N., Worthington-White, D., Garden, O. A. & Musk, P. Apoptosis of leukemic cells accompanies reduction in intracellular pH after targeted inhibition of the Na+/H+ exchanger. *Blood*, The Journal of the American Society of Hematology 95, 1427–1434 (2000).

68 Zhao, Y. et al. Targeted inhibition of MCT4 disrupts intracellular pH homeostasis and confers self-regulated apoptosis on hepatocellular carcinoma. Experimental Cell Research 384, 111591 (2019).

69 Che, X.-F., Zheng, C.-L., Akiyama, S.-I. & Tomoda, A. 2-Aminophenoxazine-3-one and 2-amino-4, 4α-dihydro-4α, 7-dimethyl-3H-phenoxazine-3-one cause cellular apoptosis by reducing higher intracellular pH in cancer cells. Proceedings of the Japan Academy, Series B 87, 199–213 (2011).

70 Hyun, S. Y. et al. Induction of apoptosis and differentiation by Na/H exchanger 1 modulation in acute myeloid leukemia cells. Biochemical and biophysical research communications 519, 887–893 (2019).

71 Sanhueza, C. et al. Sodium/proton exchanger isoform 1 regulates intracellular pH and cell proliferation in human ovarian cancer. Biochimica et Biophysica Acta (BBA)-Molecular Basis of Disease 1863, 81–91 (2017).

72 Kovacs, G. G. et al. Changes in intracellular Ca2+ and pH in response to thapsigargin in human glioblastoma cells and normal astrocytes. American Journal of Physiology-Cell Physiology 289, C361–C371 (2005).

73 Man, C. H. et al. A novel tescalcin-sodium/hydrogen exchange axis underlying sorafenib resistance in FLT3-ITD+ AML. *Blood*, The Journal of the American Society of Hematology 123, 2530–2539 (2014).

