## Supplementary Materials for "Stress pathway outputs are encoded by pH-dependent phase separation of its components"

Figs. S1 to S4

Tables S1 to S4

References (58-73)

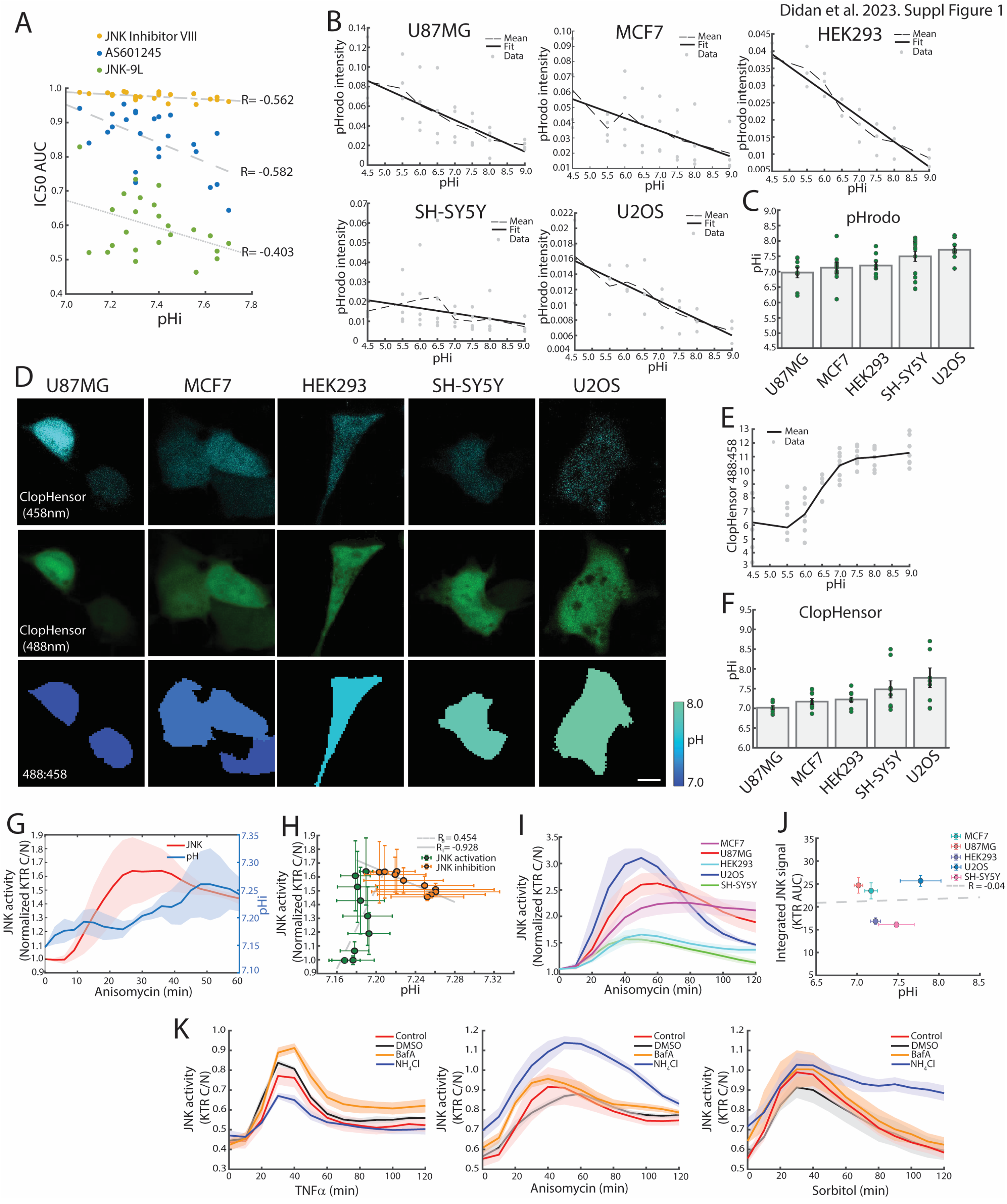


**Fig. S1. Cell lines differ in intracellular pH and response to stress treatment.** (**A**) Correlation between intracellular pH (pHi) and efficiency of anticancer drugs targeting JNK activity expressed as area under curve (AUC) of viability drug $IC_{50}$ plots. (**B**) Calibration curves for pHrodo staining in U87MG, MCF7, HEK293, SH-SY5Y and U2OS cell lines (n ≥ 8, per condition). (**C**) pHi in different cell lines measured with pHrodo. (**D**) Representative images of ClopHensor expression in different cell lines. (**E**) Calibration curve for ClopHensor measured pH in HEK293 cell line. (**F**) pHi in different cell lines measured with ClopHensor (n = 8, per condition). (**G**) JNK activity and pHi during time course treatment with anisomycin (10 ng/ml, n = 5) and (**H**) correlation between JNK activity and pHi during activation and inactivation phase. (**I**) Sorbitol-stimulated JNK activation across a panel of cell lines, n ≥ 3 independent experiments. (**J**) Association between pHi and sorbitol-stimulated JNK signal capacity measured as integrated area under JNK-KTR C/N curve across cell types. (**K**) Treatment with 50 mM $NH_4Cl$ and 50 µM Bafilomycin A1 modulates JNK activation in response to treatment with 10 ng/ml TNFα, 10 ng/ml anisomycin, and 150 mM sorbitol in HEK293 cells ($n \geq 3$, per condition). Values represent mean±SEM. "R" indicates Pearson correlation coefficients between JNK activity and pHi including correlation coefficients between JNK activation ($R_a$) or inactivation ($R_i$) and pHi. Scale bars, 10 µm.

### Figure S2

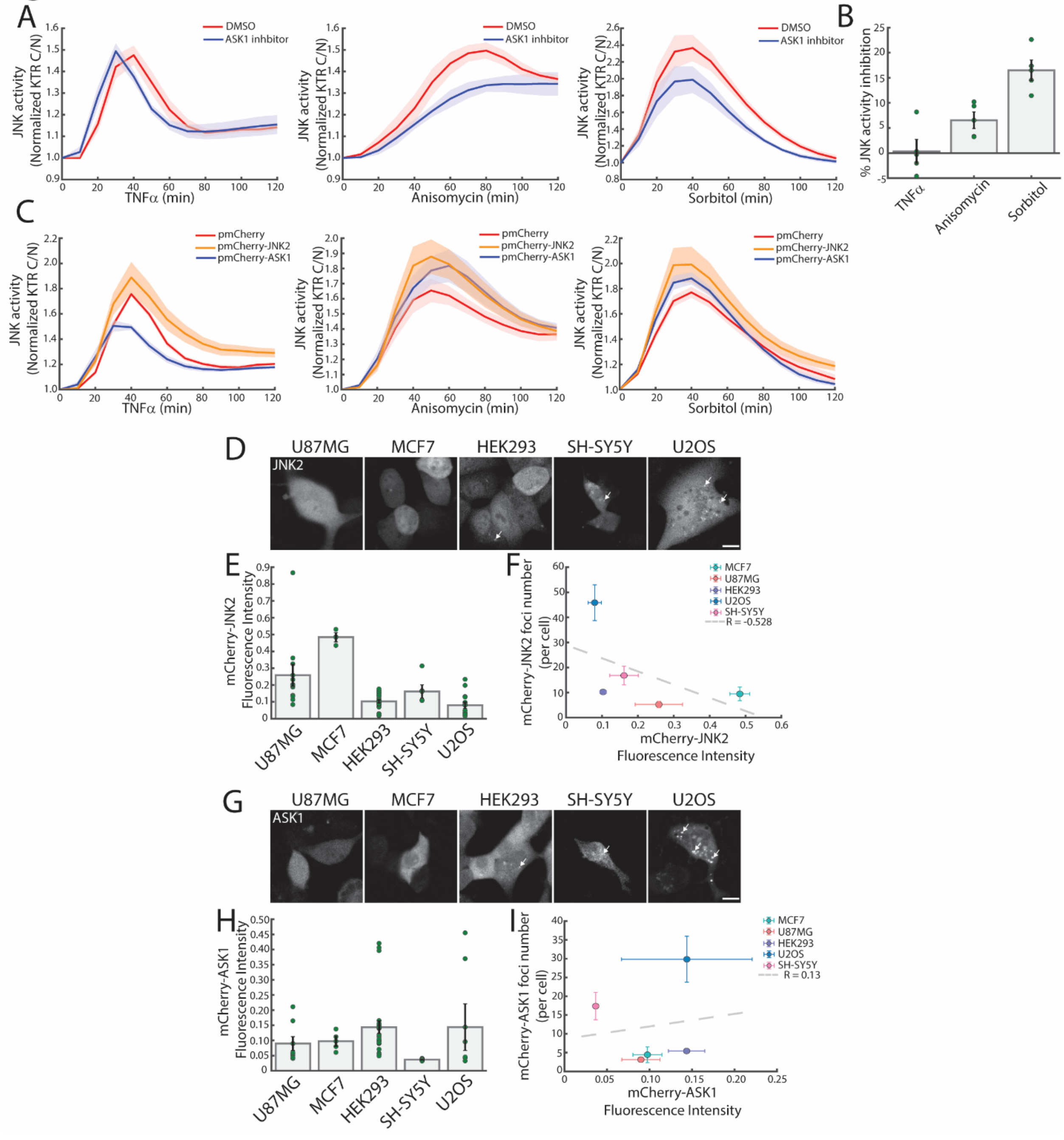


**Fig. S2. JNK2 and ASK1 transduction of stress signals and ectopic expression in mammalian cells.** (**A**) Impact of 10 µM imidazopyridine ASK1 inhibitor on TNFα (10 ng/ml), anisomycin (10 ng/ml) and sorbitol (150 mM) stimulated JNK activity (KTR), n = 4. (**B**) Quantified decrease in JNK activity (integrated area under JNK-KTR C/N curve) in presence of ASK1 inhibitor, n = 4. (**C**) Effect of ectopic expression of mCherry-tagged JNK2 and ASK1 on TNFα (10 ng/ml), anisomycin (10 ng/ml) and sorbitol (150 mM) stimulated JNK activity (KTR), ($n \geq 4$, per condition). (**D**) Transient expression of mCherry-JNK2 in cultured cell lines. (**E**) mCherry-JNK2 fluorescence intensity across a panel of cell lines ($n \geq 3$, per condition). (**F**) Correlation between mCherry-JNK2 fluorescence intensity and foci number. (**G**) Transient expression of mCherry-ASK1 in cell line panel. (**H**) mCherry-ASK1 fluorescence intensity across a panel of cell lines, ($n \geq 4$, per condition). (**I**) Correlation between mCherry-ASK1 fluorescence intensity and foci number. Values represent mean±SEM. “R” indicates Pearson correlation coefficients between pHi and JNK activity. Scale bars, 10 µm.

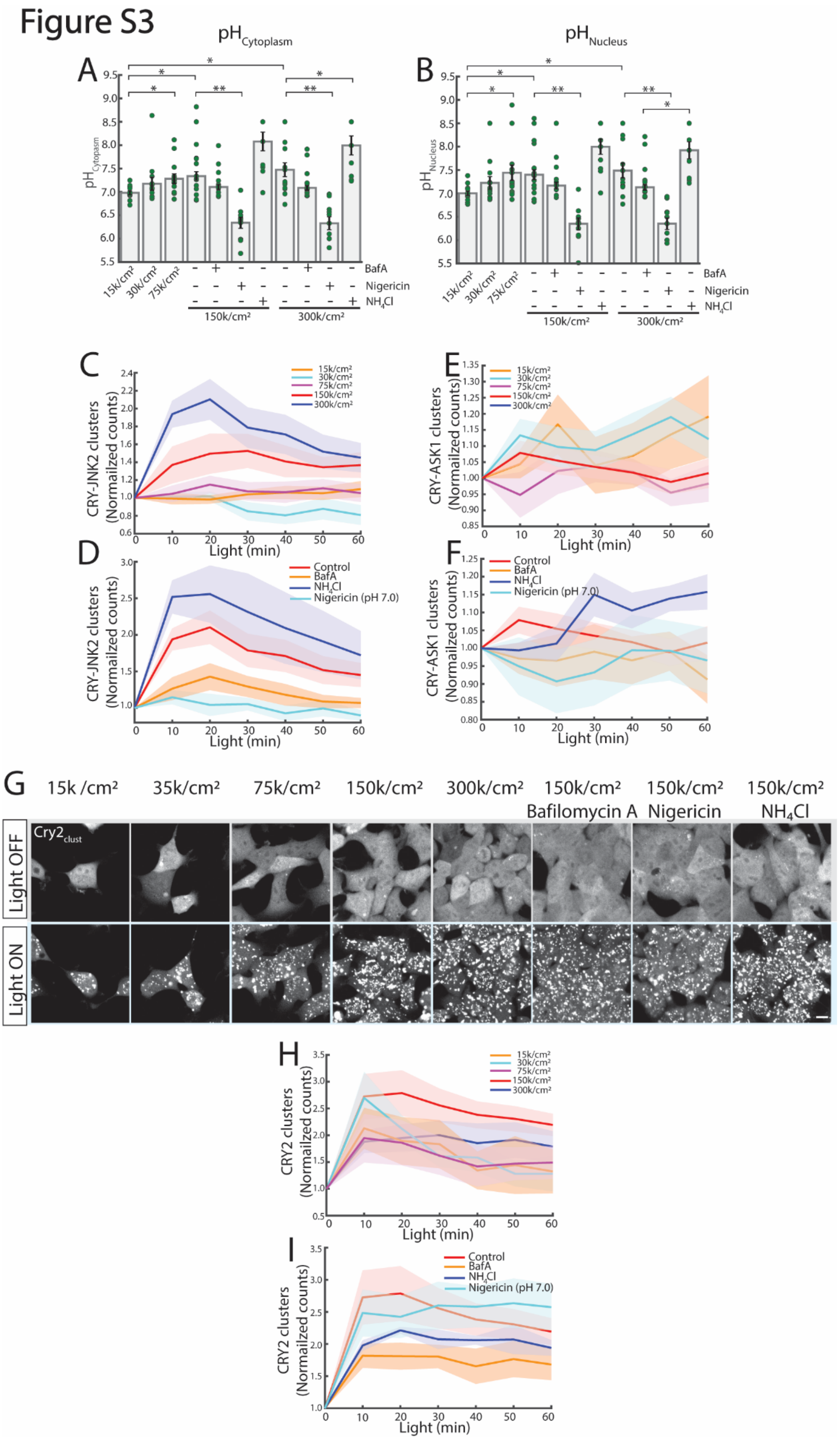


**Fig. S3. Light induced formation of JNK kinase clusters is regulated by pHi.** (**A**) Impact of cell density and chemical modulators on cytoplasmic and (**B**) nuclear pH (ClopHensor) in

HEK293 cells ($n \geq 8$, per condition). (**C**) Quantification of light induced CRY-JNK2 cluster counts in HEK293 cells seeded at different densities or (**D**) with pHi modifying chemical treatments Bafilomycin A1 (50 μM), nigericin (10 μM to equilibrate to pH 7.0) or $NH_4Cl$ (50 mM), ($n \geq 4$, per condition). (**E**) Light induced CRY-ASK1 cluster counts in HEK293 cells seeded at different densities or (**F**) with pHi modifying chemical treatments, ($n \geq 4$, per condition). (**G**) $Cry2_{clust}$ in HEK293 cells seeded at the indicated cell densities were pretreated with Bafilomycin A1 (50 μM), nigericin (10 μM to equilibrate to pH 7.0), $NH_4Cl$ (50 mM) or left untreated and light stimulated (Light On, 30 min). (**H**) Quantification of light-stimulated $Cry2_{clust}$ numbers in HEK293 cells seeded at different densities or (**I**) in presence of pHi modifying chemical treatments ($n \geq 3$, per condition). Values represent mean±SEM. Unpaired t test, *$p < 0.05$, **$p < 0.001$. Scale bars, 10 μm.

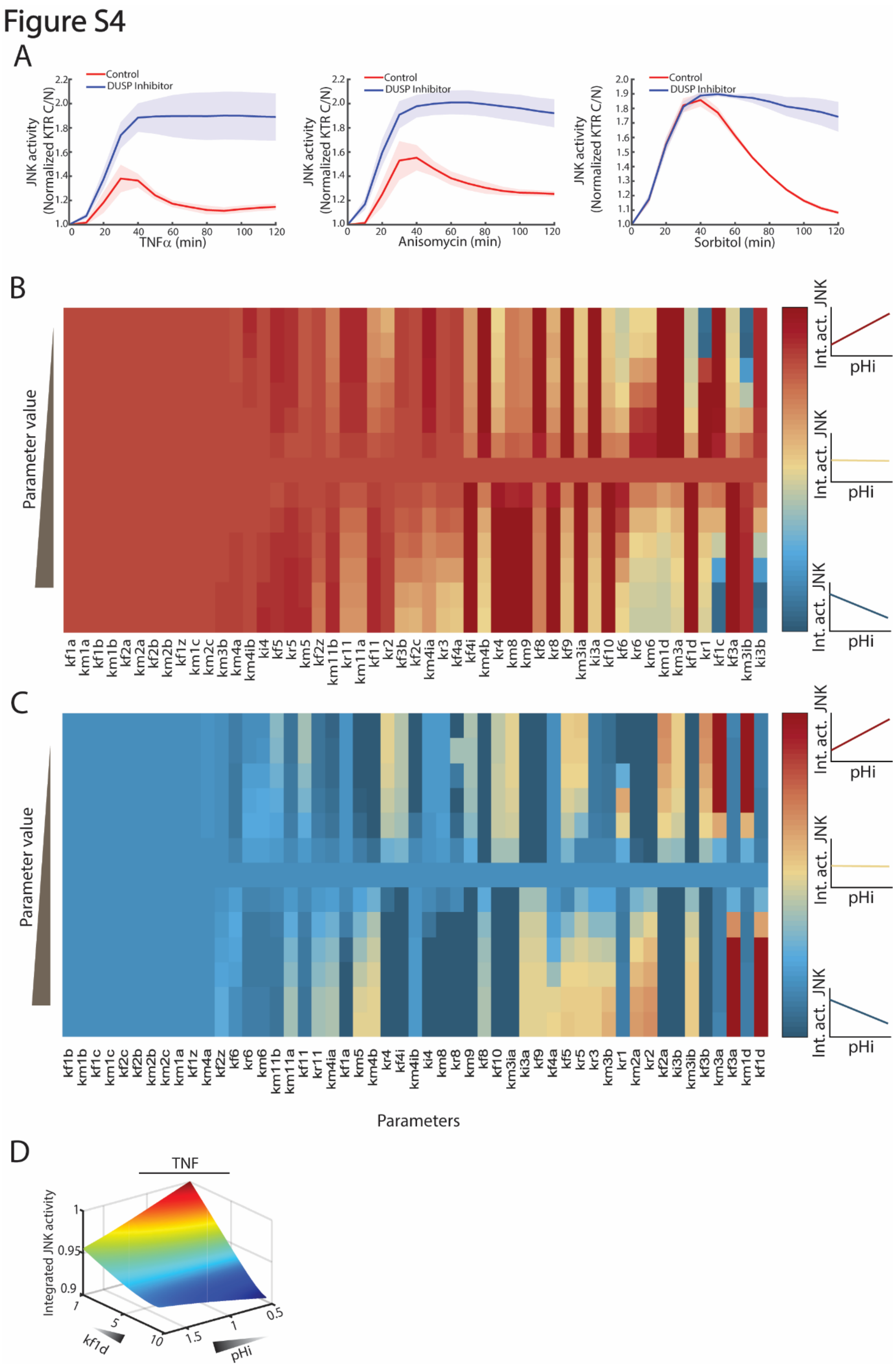


**Fig. S4. Negative feedback of JNK signals by DUSP1 and model-based sensitivity analysis of the pHi-dependent integrated JNK activity.** (**A**) Pretreatment with DUSP1 inhibitor (BCI,

10 μM) enhanced JNK activity response to TNFα (10 ng/ml), anisomycin (10 ng/ml), and sorbitol (150 mM) treatment (n = 5, per condition). Values represent mean±SEM. (**B** and **C**) Model-based sensitivity analysis, where each kinetic parameter of our model was systematically perturbed (1000-fold up or down from the nominal values), and the resulting effects on integrated JNK activity were computed. Red, blue, and yellow indicate the parameter values that cause an increase, decrease, or no change in integrated JNK activity in response to pHi concentration, respectively. (**D**) TNFα-stimulated JNK activity levels are dependent on pHi and *kf1d* parameter (ASK activation rate by ASKcon). At low *kf1d* values, there is a negative correlation between pHi levels and JNK activity. As *kf1d i*ncreases, the correlation between pHi levels and JNK activity becomes positive.

### Table S1.

**Intracellular pH of cell lines**

| Cell lines | pHi measurement method | pHi | References |
|---|---|---|---|
| K-562 | SNARF | 7.06 | [58] |
| HT-29 | SNARF-1 | 7.1 | [59] |
| MDA-MB-468 | SNARF-1 | 7.2 | [60] |
| RT-112 | SNARF1 | 7.2 | [61] |
| RPMI-8226 | BCECF | 7.25 | [62] |
| HL-60 | SNARF | 7.26 | [58] |
| U-266 | BCECF | 7.3 | [62] |
| LoVo | BCECF | 7.3 | [63] |
| 786-0 | SNARF | 7.3 | [64] |
| U-87-MG | BCECF | 7.3 | [65] |
| MDA-MB-231 | SNARF-1 | 7.32 | [66] |
| U251 | BCECF | 7.38 | [65] |
| THP-1 | SNARE | 7.4 | [67] |
| MOLT-4 | SNARE | 7.4 | [67] |
| HeLa | SNARF | 7.4 | [64] |
| Hep3B2-1-7 | pHrodo, BCECF | 7.44 | [68] |
| A431 | BCECF | 7.56 | [69] |
| KG-1 | BCECF | 7.45 | [70] |
| ACHN | BCECF | 7.65 | [69] |
| OCI-AML2 | BCECF | 7.55 | [70] |
| CCRF-CEM | SNARF | 7.18 | [58] |
| A2780 | BCECF | 7.621 | [71] |
| SK-MG-1 | BCECF | 7.65 | [72] |
| MV-4-11 | SNARF-1 | 7.7 | [73] |

**Table S2.**

**Oligonucleotides used in the study**

| Name | Oligonucleotide |
|---|---|
| NotI_iRFP F | ATTGCGGCCGCATGGCTGAAGGCTCCGTC |
| SalI_iRFP R.1 | CGGGTCGACTTACTCTTCCATCACGCCGATCT |
| BsrGI_JNK2alpha2 F | ATTTGTACACCATGAGCGACAGTAAATGTGACAGTC |
| JNK2_Hpa R | CGGGTTAACTCATCGACAGCCTTCAAGGGGTC |
| BspEI_JNK2alpha2 R.1 | CGGTCCGGATTCGACAGCCTTCAAGGGGTC |
| ASK_SgrAI_F | ATTCACCGGCGATATGAAGGCGAGCTGCCGC |
| HpaI_ASK1 R | CCGGTTAACAGTCTGTTTGTTTCGAAAGTCAATGAT |
| SmaI_ASK1 F.1 | ATTCCCGGGATATGAAGGCGAGCTGCCGC |
| SpeI_Cry2 F | ATTACTAGTAGTTCCGCGTTACATAACTT |
| PmeI_CRY2 R | CGGGTTTAAACTTATGTTTCAGGTTCAGGGG |
| PmeI_ASK1 R.1 | CGGGTTTAAACTCAAGTCTGTTTGTTTCGAAAGTCAA |
| JNK_del_388_400_F | ACTGAGCAGACGCTGGCC |
| JNK_del_388-400_R | AGGAGTGGCGTTGCTACTTAC |
| JNK_del_404-417_F | GGACCCCTTGAAGGCTGT |
| JNK_del_404-417_R | CTGCTCAGTGGACATGGATG |
| JNK_del_388-417_R | AGGAGTGGCGTTGCTACTTA |
| ASK_del_563-581_F | CAGAGCTACTGGGAAGTTG |
| ASK_del_563-581_R | CCGGAGCTCAAAGGAAGA |
| ASK_del_1291-1304_F | CATGATTCCCAGAGTGCTC |
| ASK_del_1291-1304_R | AGCATCTTCAATGACAGC |
| ASK_del_178-202_F | GCATATGTGATCAACGAAGCG |
| ASK_del_178-202_R | CGGACAACCGATGCCAGG |

Restriction site
Insertion
Gene sequence
Non target sequence

F - forward primers
R - reverse primers

**Table S3.**

**Generated cell lines**

| **Name** | **Expressed proteins** | **Antibiotic resistance** |
|---|---|---|
| U87MG_JNK-KTR | JNK-KTR-iRFP713 | Blasticidin |
| MCF7_JNK-KTR | JNK-KTR-iRFP713 | Blasticidin |
| HEK293_JNK-KTR | JNK-KTR-iRFP713 | Blasticidin |
| SH-SY5Y_JNK-KTR | JNK-KTR-iRFP713 | Blasticidin |
| U2OS_JNK-KTR | JNK-KTR-iRFP713 | Blasticidin |
| HEK293_ClopHensor | JNK-KTR-iRFP713, ClopHensor | Blasticidin, G418 |
| HEK293_JNK2 | JNK-KTR-iRFP713, JNK2-mCherry | Blasticidin, G418 |
| HEK293_ASK1 | JNK-KTR-iRFP713, ASK1-mCherry | Blasticidin, G418 |
| HEK293_Cry2JNK2 | JNK-KTR-iRFP713, Cry2clust-JNK2-mCherry | Blasticidin, G418 |
| HEK293_Cry2ASK1 | JNK-KTR-iRFP713, Cryclust-ASK1-mCherry | Blasticidin, G418 |

### Table S4.

**Reactions and rate equations for the models.**

| Reaction Number | Reaction | Rate equation |
|---|---|---|
| R1 | ASK => act_ASK | kf1z + kf1a * TNF * ASK/(km1a + ASK) + kf1b * Anisomycin * ASK/(km1b + ASK) + kf1c * Sorbitol* ASK/(km1c + ASK) + kf1d * act_ASKcon * ASK/(km1d + ASK) - kr1 * act_ASK |
| R2 | MAP3K => act_MAP3K | kf2z + kf2a * TNF * MAP3K /(km2a + MAP3K)+ kf2b * Anisomycin * MAP3K /(km2b + MAP3K) + kf2c * Sorbitol* MAP3K /(km2c + MAP3K) -kr2 * act_MAP3K |
| R3 | JNK => act_JNK | kf3a * act_ASK * JNK/(km3a+ JNK)+ kf3b * act_MAP3K * JNK/(km3b+ JNK) - ki3a * DUSP1 * act_JNK/(km3ia + act_JNK) - ki3b * JNK2con * act_JNK/(km3ib + act_JNK) - kr3 * act_JNK |
| R4 | => pHi | (kf4base + kf4a * act_ASK/(km4a+ act_ASK)) - (kf4i * activepump/(km4ia + activepump)) - ki4 * act_JNK * pHi /(km4ib+ pHi) - kr4 * pHi/(km4b+pHi); |
| R5 | JNK2 => JNK2con | kf5 * pHi * JNK2/(km5+ JNK2) - kr5 * JNK2con |
| R6 | nASKcon=> act_ASKcon | kf7 * pmAkt309 * amTorc2 / (Km8 + pmAkt309) - kr7 * pmAkt309474 |
| R7 | => RNADUSP1 | kf8 * act_JNK /(km8+ act_JNK) - kr8 * RNADUSP1 |
| R8 | RNADUSP1 => DUSP1 | kf9 * RNADUSP1 /(km9+ RNADUSP1) |
| R9 | DUSP1 => | kf10 * DUSP1 |
| R10 | pump => activepump | kf11 * pHi * pump/(pump+km11a) - kr11 * activepump/(activepump+km11b) |
